# Phospholipid signaling and multivesicular endolysosomes modulate membrane dynamics at the biotrophic interface in Rice Blast

**DOI:** 10.1101/2024.11.04.622007

**Authors:** Poonguzhali Selvaraj, Fan Yang, Wenhui Zheng, Naweed I. Naqvi

## Abstract

*Magnaporthe oryzae*, the rice blast pathogen, secretes an arsenal of apoplastic and cytoplasmic effector proteins to suppress host immunity during the biotrophy phase. The proposed non-conventional secretion of cytoplasmic effectors into the Biotrophic Interfacial Complex (BIC), remains uncharted except for the involvement of the exocyst complex therein. Recently, the plant endocytic machinery has been implicated in translocation of cytoplasmic effectors to the host. Here, we used live cell imaging and mutant analyses to provide new insight into late endolysosomal mediators as conduits for establishment of the BIC and/or unconventional secretion across the host interface. Rerouting of the late endocytic GTPase Rab7 occurs at the BIC; and the dominant negative Rab7^T22N^ led to significant defects in focal interface formation. FYVE biosensor and pharmacological interventions revealed that Rab7 membrane trafficking and establishment/integrity of the biotrophy interface requires the PI3 kinase Vps34 and phosphoinositide PI3(P) function. Additional pharmacological evidence supports endolysosomal trafficking to the host interface follows a distinct non-conventional secretion route too. Finally, we show that the Multi-Vesicular Bodies and lysosomal membranes act as downstream modules of the Rab7 function in precise sorting and specialised traffic of cytoplasmic effectors across the *Magnaporthe*-Rice interface during blast disease.

## Introduction

Filamentous plant pathogens such as fungi and oomycetes orchestrate elaborate manipulation of host cells to accommodate infection structures that frequently require recruitment of host-derived membranes and formation of novel interfacial structures. These peri-microbial interfaces are critical sites that facilitate nutrient uptake and prolonged defence evasion. The biotrophic interfacial complex (BIC), a novel, specialized interface produced by the devastating rice-blast pathogen *Magnaporthe oryzae*, represents one such focal membranes and cytoplasm-rich structure in host plants. *M. oryzae* is an aggressive pathogen that infects other cereals and millets and more recently a notorious pathotypic variant of this genus has emerged to cause wheat blast in South America, Bangladesh and Zambia (Arora et al., 2021). Blast disease continues to pose a major threat to global food security affecting wheat and rice production worldwide (Garcia et al., 2023). The Ascomycete pathogen uses a multistage hemibiotrophic infection strategy: dome-shaped appressorium uses a brute-force attack mechanism to puncture the host surface and form penetration peg that transits to tubular filamentous primary invasive hyphae (IH) first and later to the bulbous biotrophic intracellular invasive hyphae developing inside living epidermal cells, and remain enveloped by an intact host plasma membrane termed the extra-invasive hyphal membrane (EIHM) and finally switches to the necrotrophic killing phase within the host tissues. The BIC is formed in each newly infected cell at the tip of the primary IH and later positioned subapically as the bulbous IH develops and serves as the primary site for deployment of discrete effector proteins.

The repertoire of *M. oryzae* effector proteins secreted to thwart the activation and execution of plant defense responses is classified based on the origin of secretion and destination in the interaction milieu. The apoplastic effectors, governed by the conventional ER-to-Golgi secretory process sensitive to Brefeldin A (BFA), remain secured to the EIHM inside the host, while the cytoplasmic effectors destined for translocation into host cells via the BIC are secreted through an unconventional BFA-insensitive pathway involving the exocyst and SNAREs (Giraldo et al., 2013, Bin Qian et al., 2022). Despite major advances in effector biology and the pivotal role of BIC in effector delivery, remarkably little is known about the biogenesis of this interfacial zone and the contribution made by the fungus, if any, to this interface remains elusive. Alternatively, rapidly expanded research on haustoria, the interfacial hyphal structures formed by Oomycetes and other filamentous pathogens revealed that rerouting of the host endocytic components and diversion of autophagy machinery at the haustorial interface (Bozkurt and Kamoun, 2020; Rausche et al., 2020, Peter et al., 2021) indicating massive reprogramming of host membrane trafficking during infection. Recent evidence from plants further implicates extracellular vesicles or multivesicular bodies (MVB) at the haustorial interface in cross-kingdom communication during host-pathogen interaction (Cai et al., 2018; Rutter and Innes, 2018; Baldrich et al., 2019). The MVB or multivesicular bodies are a special form of the late endosomes generated when the limiting membrane of the endosome invaginates and buds into its own lumen. During this process, proteins residing in the endosomal membrane are sorted into the forming vesicles. MVB of the endocytic process are not always destined for degradation within vacuoles and can follow a retrograde pathway involving the fusion with the plasma membrane, thereby delivering their Intralumenal Vesicles (ILV) to the cell exterior and such Extracellular Vesicles (EVs) are often termed as exosomes (Schorey and Bhatnagar, 2008; Zhang et al., 2019). The finding that both bacterial pathogens and plant can discharge EVs with immunomodulatory functions (Bahar et al., 2016; Wang et al., 2017a, b; Cai et al., 2018; Baldrich et al., 2019) has sparked renewed interest in dissecting the contents and functions of the EVs deployed at interfacial matrix.

Fungal MVBs are abundant in haustoria and putative exosome vesicles have been detected in the paramural space and EHMx of the powdery mildew pathogen thus suggesting the existence of an exosome-mediated secretion in *Golovinomyces orontii* interactions (Micali et al., 2011). However, investigations on EVs from fungal phytopathogens remain rudimentary and was first reported from *Alternaria infectoria*, the leaf blight pathogen (Silva et al., 2014). Subsequent studies have examined the EVs from other phytopathogens like *Zymoseptoria tritici, Fusarium oxysporum* f. sp. *vasinfectum*, *Ustilago maydis*, *F. graminearum* and more recently from *Colletotrichum higginsianum* and the proteomics of few EVs suggests the presence of effector proteins and phytotoxic compounds (Bleackley et al., 2019; Garcia-Ceron et al., 2021a, b; Hill & Solomon 2020; Rutters et al., 2022). Nevertheless, despite a head start in the last decade, research into the role of fungal EVs in plant-pathogen interactions remains languish and the biogenesis of BICs still remain in its infancy that might partly be due to technical difficulties associated with isolating BIC complexes for analysis. BIC associated hyphal cells are rich in secretion machinery components – the exocyst and SNARE proteins (Khang et al., 2010), the docking subunits tethering vesicles to the secretory sites (Heider & Munson, 2012). The Qc-SNARE protein MoSyn8 and the endocytic protein MoEnd3 are involved in the secretion of cytoplasmic effectors while dispensable for apoplastic effector secretion (Qi et al., 2016, Li et al., 2017). Furthermore, a recent investigation revealed that BIC contains the host-derived Clathrin-mediated endocytic (CME) components and silencing of the rice CME prevents *M. oryzae* infection (Garcia et al., 2023) which indicates that rice blast likely triggers reprogramming of the host endocytic machinery at the interface. Although, these studies indicate the significance of vesicle trafficking in rice-blast interactions, from a cell biology perspective, perhaps the most significant questions of general importance for the blast pathogen remain to be explored. How do the effectors get secreted by invasive hyphae and route in to the BIC interface? Does the fungus use exocytic mechanisms for the generation of exosomes like EVs as reported to occur in some plant-fungus interactions and if so how are they formed and how do they route into the host cells interface with plant endocytic mechanism? (Xia-Yan and Talbot, 2016).

To answer such questions, we set out to investigate the role of vesicular trafficking in effector secretion in *M. oryzae* and further assessed if a fight back mechanism by the pathogen to the modulation of endocytic machinery of the host components at the BIC interface. The specificity of vesicular transport relies mainly on a key component the Rab GTPases that acts as molecular switch cycling between active and inactive states and co-ordinate the machineries for endosomal maturation, trafficking and membrane fusion (Uemura and Ueda, 2014). Rab7-GTPase regulates the transport and maturation of acidic late endosomes as well as their fusion with lysosomes/vacuoles thus regulate transition of late endosomes into MVBs and play a fundamental role in controlling late endocytic membrane traffic. Pathogen-secreted effector proteins have emerged as robust molecular probes and can be leveraged to label specific subcellular compartments to reveal novel and important processes in host cells (Bozkurt et al., 2015, Toruno et al., 2016; Peter et al., 2021). Here, we used blast effectors and Rab7 of *M. oryzae* as markers to study vesicle trafficking processes at the BIC interface by live cell imaging of the invasive hyphae formed during infection. We report here that in *M. oryzae*, vacuole targeted late endosome Rab7 is diverted to the BIC interface and this rerouting is specific for the secretion of cytoplasmic effectors. We confirmed the robustness of our results through mutant analysis and show that reduced pathogenicity of Rab7 mutants is due to the impairment of focal BIC formation and henceforth attenuated invasive hyphal growth. Further, we describe Vta1 (**V**ps **t**wenty **a**ssociated protein **1**) as a marker for MVBs in *M. oryzae* and through live cell imaging, gene deletion and pharmacological investigations confirm that the MVB-mediated extracellular vesicles form the unconventional secretion system for the cytoplasmic effectors in *M. oryzae* / rice blast pathosystem.

## Results

### Rab7 marks the late endocytic events in *M. oryzae*

To decipher the function of membrane trafficking in biotrophic interface formation, we first identified a bona fide marker for the late endosomal compartments in *M. oryzae*. In yeasts and mammalian cells, Rab7 plays a fundamental role in controlling the endocytic membrane traffic at the level of late endosomes by regulating the tethering and fusion of transport vesicles and/or the lysosomes. We previously reported that the Rab7 homolog Ypt7 (mentioned as Rab7 hereafter) in *M. oryzae* localized to the late endosomal compartments and is designated as a late endosomal marker (Ramanujam et al., 2013). Through live imaging and time-lapse analyses of GFP-Rab7, we confirmed that Rab7 marks the late endocytic events during the pathogenic phase of *M. oryzae* (**Figure 1**). Throughout the pathogenic phase of *M. oryzae*, Rab7 predominantly localized to the cytoplasmic vesicles and the vacuolar membrane, as confirmed by colocalization with CMAC staining that marks the vacuoles (**Figures 1A, 1B**). Further, Rab7 marked the membrane vesicle budding, the movement and fusion of transport vesicles to the target membranes followed by fusion of these membranes (**Figures 1C-F**, Videos S1, S2 and S3), the archetypal sequential steps in the late endocytic events of vesicular membrane trafficking (Inada et al., 2016). The fluorescent vesicles were mobile and moved along the tubules bidirectionally thus facilitating the transport as new vesicles pinched off from the membranes. Nevertheless, Rab7 marked the MVBs - the late endosomes with intra luminal vesicular structures in conidia and the invasive hyphae (**Figure 1**). These results confirmed that Rab7 marks the late endocytic and endolysosomal events in *M. oryzae* and represents an ideal marker to visualize the vesicular trafficking at the pathogen-host interface and to assess if MVB-mediated extracellular vesicles are involved in cytoplasmic effector secretion in rice blast.

**Figure 1.**
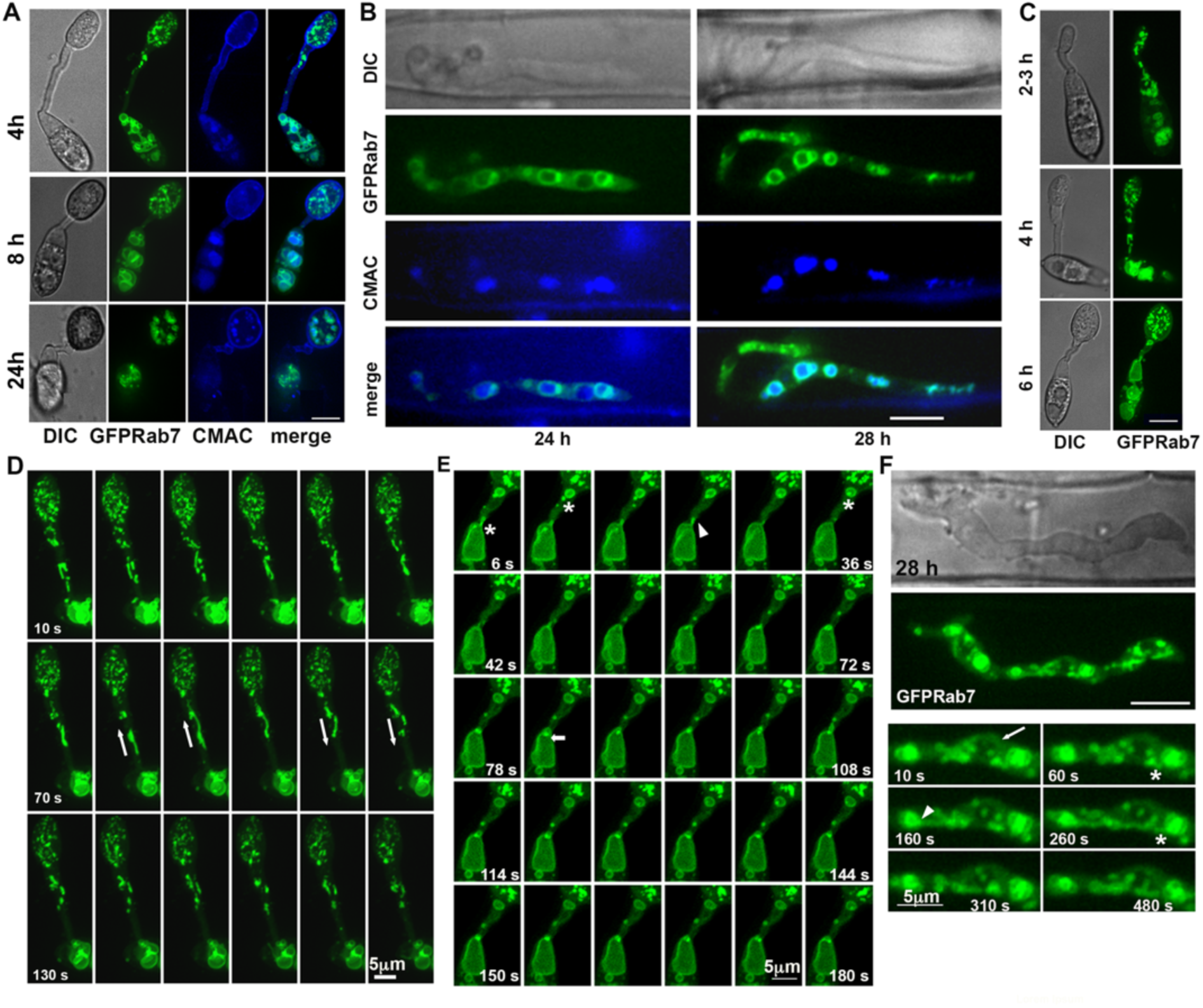
Rab7 marks the late endocytic and endolysosomal membrane compartments in *M. oryzae*. (A-C) Localization of GFP-Rab7 at different stages of infection in *M. oryzae*. (A) Co-localization of GFP-Rab7 CMAC in the infection-structures (A) and the invasive hyphae (B). The images shown are maximum projection of 5 planes at 0.3 μm spacing. (D,E) A montage of the time lapse analysis obtained at 4 h (D) and 6 h (E) is shown. The GFP-Rab7 punctae or vesicles traverse from the conidia to the emerging appressoria as a continuum of tubular vesicles (white asterisks) sometimes enlarging to long tubular vacuoles (white arrows) via fusion of the smaller vesicles. (E) A prominent large vacuole appears at the terminal cell of the conidium as the appressorium matures, and a small, pointed vesicle that buds from the plasma membrane (white arrow) and pinches out of the vacuole (white arrowhead) and moves into the appressorium by fusion and fission events (white *). (F) Subcellular localization of GFP-Rab7 in the invasive hyphae *in planta* in vesicles and vacuolar membrane marking the different stages of vesicle budding (white asterisk) and vesicle fusion (white arrowhead) and tubular vacuoles (white arrows).

### Rab7 GTPase activity is crucial for focal BIC formation and pathogenicity of *M. oryzae*

Effectors locate to discrete locations in the invasive hyphae based on the secretion mechanism and have been visualised in real time by live cell imaging in *M. oryzae* (Khang et al., 2010). To investigate the role of such late endocytic and vacuolar machinery in BIC formation and/or secretion of effectors, we epifluorescently labelled Pwl2 (Pwl2-mCherry) or Bas4 (Bas4-GFP or Bas40-mCherry) as candidates for cytoplasmic and apoplastic effectors, respectively, and co-expressed them with GFP-Rab7 (gRab7Pwl2 or gRab7Bas4, **Table S1**) to visualize the vesicular dynamics at the fungus-host interface (**Figure S1**). The formation of BIC in the first invaded cell involves three developmental and functional stages - the tip BIC, the early side BIC and the late side BIC mentioned as focal BIC hereafter (**Figure 2A**) that remains in the first cell while invasive hyphae continue to proliferate (Jones et al., 2021). GFP-Rab7 localized to endosomal vesicles, vacuolar membrane and MVBs in the invasive hyphae. Most interestingly, a multivesicular body or vesicle remained in close proximity to and colocalized with the Pwl2-mCherry (BIC) at specific stages of biotrophy. The MVB adjacent to the BIC co-localized with Pwl2mC during the tip BIC and side BIC stages, whereas the MVB appeared to move significantly away from the BIC during the focal BIC stage compared to the previous junctures (**Figure 2B**). However, no such association between GFP-Rab7 positive structures and Bas4mC marked EIHM could be observed in *M. oryzae* (**Figure S2**). The embossment of MVB beneath the BIC and its colocalization with Pwl2mC at specific stages revealed that Rab7 is involved in *Magnaporthe* / rice interaction. Nevertheless, to gain a concrete evidence on the effect of Rab7 in focal BIC formation, we generated a dominant negative variant (Rab7^DN^ - Thr22Asn) locked in GDP conformation, and a constitutive active GTP bound (Rab7^CA^ - Glu67Leu) Rab7 mutant and investigated the biotrophy interface therein. In the wild type strain, Rab7^DN^ or Rab7^CA^ mutant version driven by the native promoter, showed growth defects in hyphal, loss of conidiation, and significantly reduced pathogenicity (**Figure S3**). Henceforth, conditional mutants that expressed amino terminal tagged GFP-Rab7 driven by the Pwl2 promoter that is expressed only during the IH growth, were generated along with the WTRab7 (PgRab7). Such conditional mutants were then co-expressed with Pwl2 (PPgRab7) or Bas4 (BPgRab7) effector fused with mCherry and investigated. Despite having no defects in hyphal growth, conidiation or appressorium formation, the mutant Rab7^DN^ had impaired invasive hyphal growth and was less pathogenic, while Rab7^CA^ was comparable to Rab7^WT^. The membrane localization of Rab7 was strictly dependent on the nucleotide switch of the GTPase, as the Rab7^DN^ was cytosolic; whereas the vesicular and membrane association of Rab7 was restored in the Rab7^CA^ strain when marked with either of the effectors as in the wild type (**Figures S4, S5, Figure 2**). In addition to impaired hyphal growth and cytosolic Rab7, the PPgRab7^DN^ mutant expressing Pwl2mC showed defective BIC formation; the tip or side BIC at earlier stages was either not evident or delayed and the focal BIC formed was either dispersed, or split or not properly located within the focal plane. The focal BIC formation or the Pwl2mC localization was unaffected in the PPgRab7^CA^ and comparable to PPgRab7 (**Figures 2c and 2d**). Rather surprisingly, though BPgRab7^DN^ marking Bas4mC had reduced pathogenicity, the secretion at EIHM or the apoplastic effector Bas4mC *per se* was not affected (Figure S5). It is known that focal BIC formation is essential for proper proliferation of the invasive hyphae and impaired BIC results in reduced pathogenicity (Nishimura et al., 2016). Nevertheless, our results confirmed that the reduced pathogenicity of Rab7^DN^ mutant due to the defect in proliferation of invasive hyphae is brought by the impaired BIC and suggest that proper spatiotemporal regulation of Rab7 GTPase activity-mediated vesicle trafficking are critical for proper focal BIC formation in *M. oryzae*

**Figure 2.**
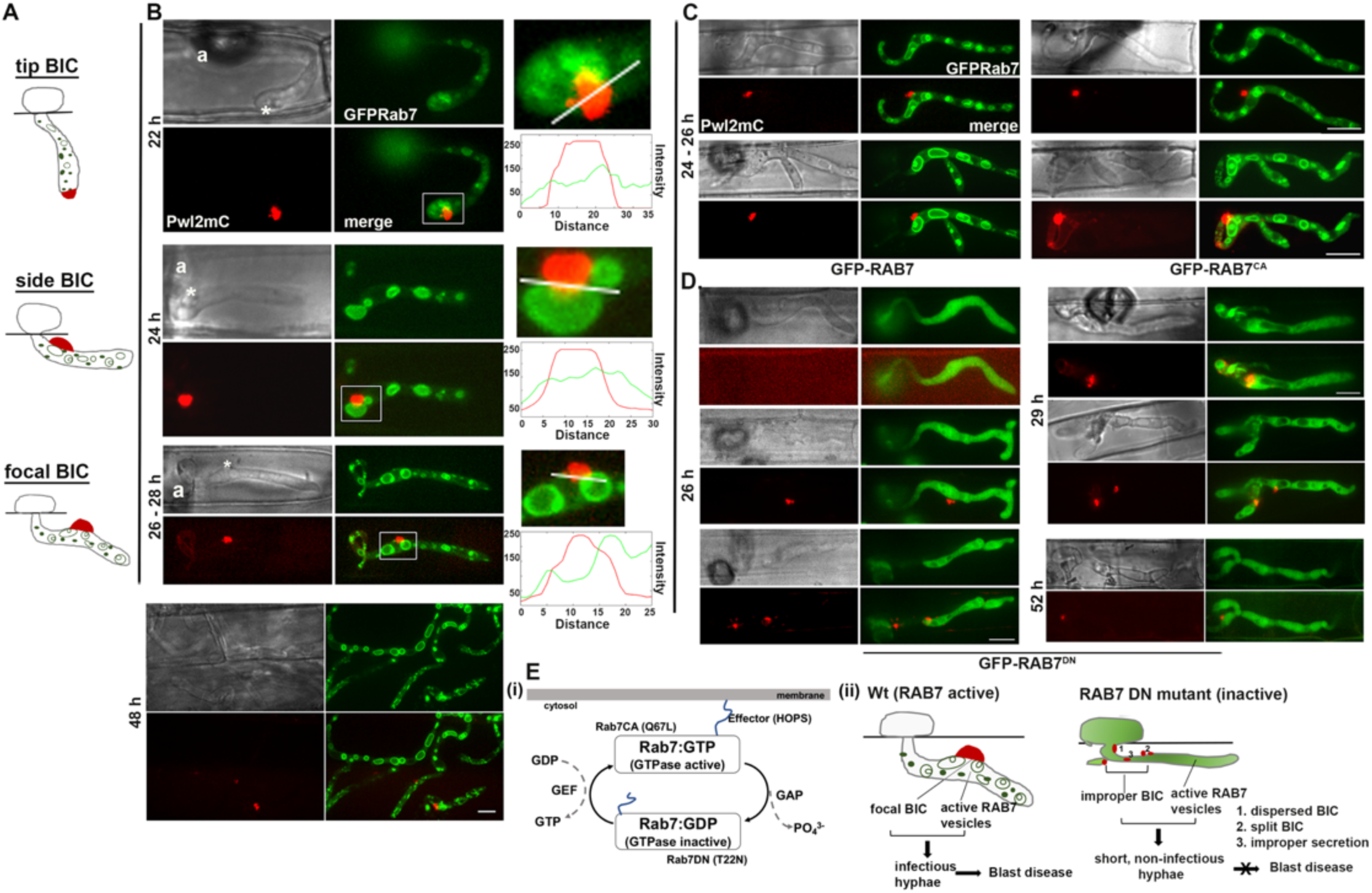
The Rab7 GTPase activity is essential for establishing the focal biotrophy interface and pathogenicity in *M. oryzae*. (A) Schematic depiction of the different stages of biotrophy interface formation in *M. oryzae*. (B) Micrographs showing the localization of GFP-Rab7 and Pwl2-mCherry in the invasive hyphae at earlier time points corresponding to BIC stages and a later time point (48 h). The insets (white rectangle) show enlarged view alongside the corresponding fluorescence intensities of the demarcated region (white line) on the right. (C) the localization of GFP-Rab7 and Pwl2-mcherry in the *PgPRab7* strain (GFP-Rab7), PgPRab7^CA^ (GFP-Rab7^CA^) and (D) PgPRab7^DN^ (GFP-Rab7^DN^). The confocal images are maximum projection of 13 z stacks of 0.3 μm width at 0.5 μm thickness. Scale bar represents 10 μm. (E) Schematic showing the Rab7 configuration (i) and the effect of Rab7^DN^ mutant on BIC formation and invasive hyphal growth (ii) GFP-Rab7 localize to late endosomes and vacuolar membrane and the BIC formation is normal in WT while the GFP-Rab7^DN^ mutant protein is sequestered in the cytosol and vesicle trafficking is perturbed and further causes deformities in BIC formation that ultimately leads to reduced pathogenicity in *M. oryzae*.

### RAB7-mediated late endocytic trafficking is rerouted and MVB interacts with BIC through the ILV contents

The shift or the difference in the localization of MVBs at different stages of BIC formation hinted at a change in trafficking of late endosomes at or near the biotrophy interface. To address if rerouting of Rab7 trafficking occurs near the BIC for effector secretion, we performed time lapse analysis to monitor the GFP-Rab7 and Pwl2-mCherry at different stages of biotrophy. The results clearly revealed that GFP-Rab7 vesicles are trafficked into the Pwl2-mCherry marked biotrophy interface (**Figure 3**, **Video S4 and S5**). To our surprise, recurrent trafficking of GFP-Rab7 in to Pwl2-marked BIC could be observed consistently during the side BIC stage indicating that such rerouting might be stage specific, although MVB could still be observed beneath the focal BIC. Nevertheless, this is highly likely since tip and early side BIC functions in effector delivery, while the late side BIC or focal BIC is considered a remnant and unable to deliver effectors during the early biotrophic phase (Jones et al., 2021). Such Rab7-positive vesicles entering into the BIC interface were also evident in PPgRab7^WT^ and PPgRab7^CA^ (**Figure S6, Video S6 and S7**) implying that active Rab7 vesicle trafficking during host invasion is critical for focal BIC formation and biotrophy of *M. oryzae*. Such punctate vesicles delivered by the protrusion of the outer membrane of the MVB beneath the BIC while the intralumenal vesicles fuse and release the punctae that either fuse with the BIC membrane (Figure 3A) or enter in to the BIC (**Figure 3B**). Furthermore, time course analysis and colocalization with CMAC staining clearly revealed that it is not the outer membrane counterpart, but the intralumenal vesicles that release a small punctae delivered to the BIC. As mentioned, this rerouting to the BIC interface occurs at specific stages, while continuous vesicular trafficking occurred throughout the hyphae wherein those vesicles fuse with the nearby vacuole or the MVB membrane and thus couldn’t be imaged as separate punctae thereafter (**Figures 3C, 3D, Videos S8 and S9**). However, the exact mechanism of such rerouting of the intralumenal vesicular contents of the MVBs or late endosomes to the complex interface remains to be unravelled. Nevertheless, we used a specific probe to monitor the multi vesicular endosomes to confirm the identity of ILVs that are trafficked to the biotrophy interface,

Phosphatidylinositol 3-phosphate (PtdIns3P or PI3P) is a master regulator controlling the fundamental aspects of membrane dynamics, protein sorting and lipid signaling. PtdIns3P is mostly found on the limiting and intraluminal endosomal or endolysosomal membranes. PI3P functions by recruiting a variety of effectors, containing the PI3P-binding module FYVE or PX, that control diverse membrane trafficking events including but not restricted to inward vesiculation, and ESCRT-dependent intraluminal cargo sorting in MVBs. Therefore, we used 2×FYVE-GFP as a specific probe (Gilloly et al., 2000) to monitor PI3P on the intraluminal vesicles and the major multivesicular endosomes underneath the biotrophy interface. FYVE-GFP localized to vesicles and membranes of different sized vesicles in the infection structure and in the *in planta* invasive hyphae. The FYVE vesicles near the focal BIC partly colocalized with the Pwl2-mCherry and time lapse imaging revealed that FYVE vesicles enter the BIC (Figures 4A-C, Videos S10 and S11). Treatment with LY294002 that inhibits the PI3-kinase (Vps34) in micromolar range (Lu et al., 2013), abolished such FYVE vesicles and the GFP was retained in the enlarged vacuoles both in the conidia and in invasive hyphae (Figures 4A, 4D). Treatment with PI3-Kinase inhibitor resulted in enlarged vacuolar structures that completely lack the internal multivesicular bodies (Fernandez-Borja *et al*., 1999). Addition of LY294002 before penetration into the rice leaf sheath significantly delayed the host penetration and further impaired the overall invasive growth *in planta*. More interestingly, treatment with LY294002 led to the formation of aberrant focal BIC structures which appeared to be either abnormal, dispersed or split in two. These results helped us conclude that proper spatiotemporal regulation of the PI3-kinase activity and the MVB-mediated vesicular membrane trafficking are necessary for establishment of the focal biotrophy interface and the associated secretion machinery in rice blast.

**Figure 3.**
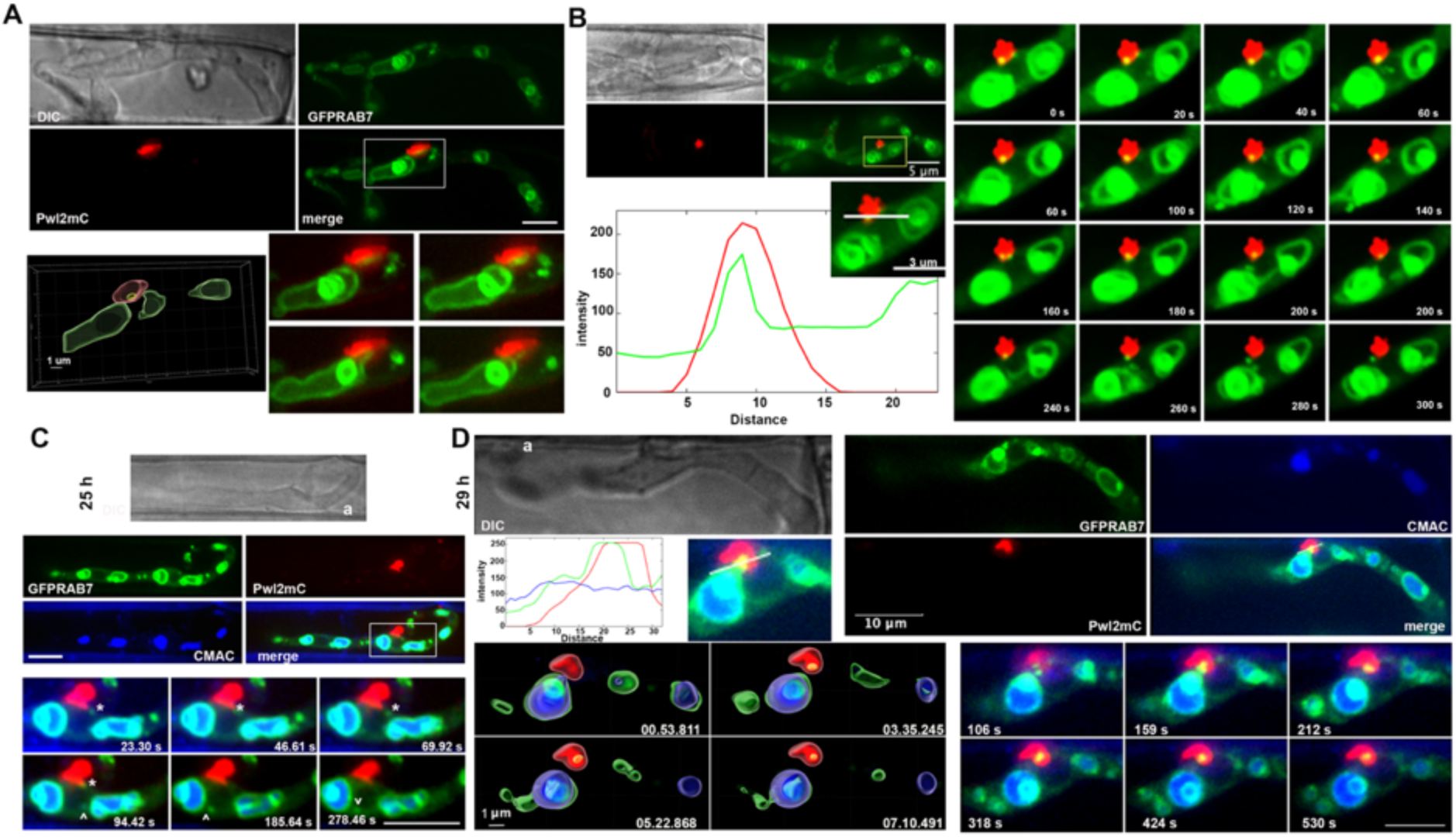
Rerouting of Rab7 trafficking at the BIC interface, and the release of intralumenal vesicles into the BIC. (A-B) Images showing the localization of GFP-Rab7 and Pwl2-mCherry in the biotrophic hyphae at the indicated timepoints in gRab7Pwl2. (A) image at the bottom right shows the selected timepoint of a sequential time lapse clearly showing the fusion of an MVB and the release of ILV into the BIC; the image at the bottom left depicts the surface rendered projection at a single timepoint wherein the MVB vesicle is evident in the BIC. (B) Panel at the left bottom shows the enlarged view of selected area (white rectangle) and the fluorescence intensity curves of the line shown therein. Right panel shows a montage of sequential images of a time lapse obtained at 20 s intervals. The images clearly show the release of a vesicular structure from the MVB that colocalizes with the Pwl2-mCherry-marked BIC. (C-D) Co-localization of GFP-Rab7 Pwl2-mCherry with the vacuole marker CMAC. (C) movement of vesicles from MVB to the biotrophy interface (white asterisks) and the conventional trafficking between the organelles (white open arrowhead). (D) CMAC staining clearly distinguishes that it is the ILV contents that get trafficked to the host interface during biotrophy (bottom right panel) and the surface rendered images for selected timepoints are shown (left bottom panel). The images shown are the z projections of a stack of 17 planes of 0.3 μm width. Scale bars unlabelled are 5 μm

**Figure 4.**
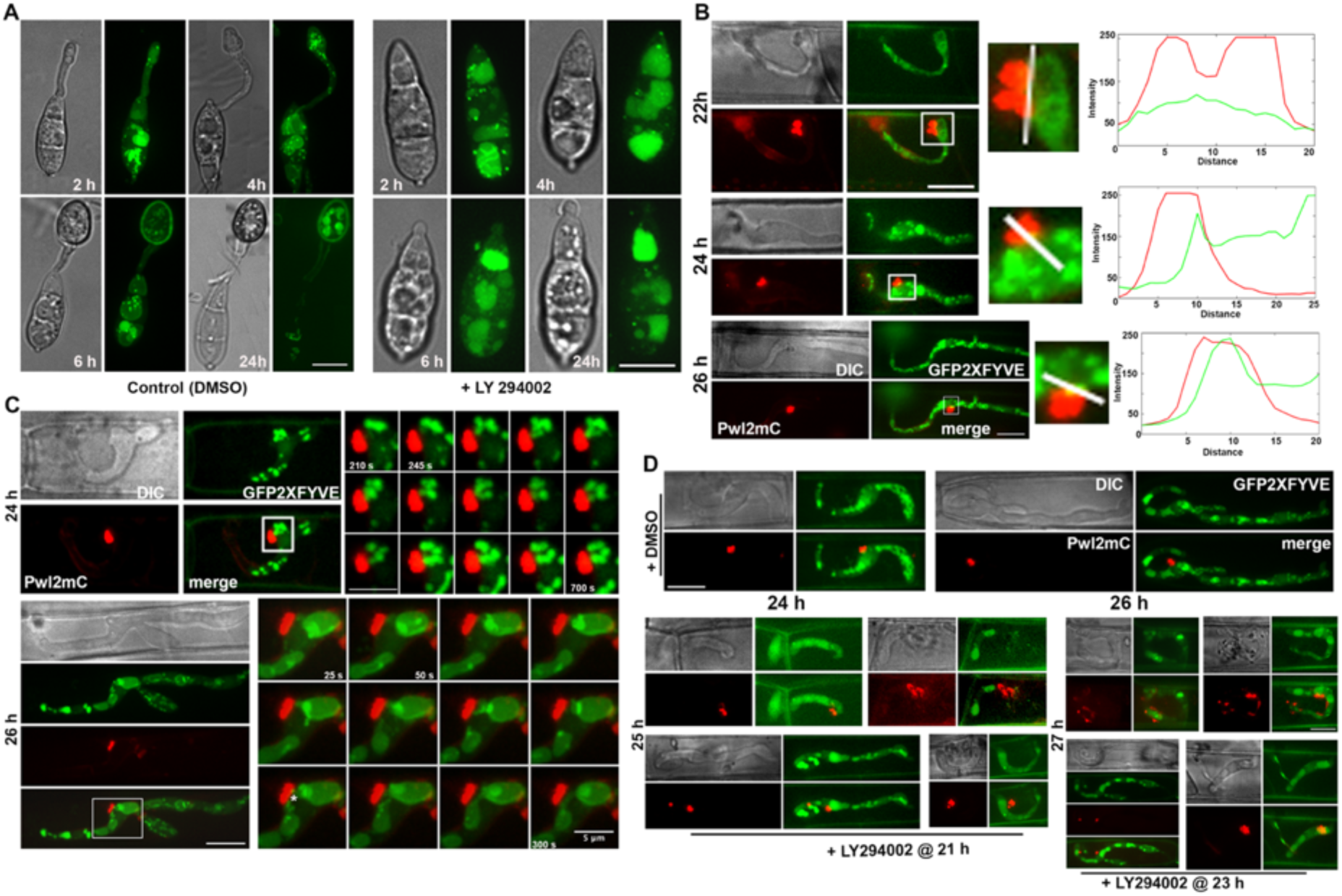
Released intralumenal vesicles are enriched in phospholipid PI3P and trafficked to the biotrophy interface. (A) Localization of GFP-FYVE vesicles in Pwl2FYVE during conidiation and appressorium formation (left) are perturbed upon treatment with the PI3P kinase inhibitor LY294002 (right). With the addition of inhibitor, FYVE membrane localization is abolished, and the number of vesicles was low, and the conidia failed to germinate and form appressoria. (B) Localization of GFP-FYVE and Pwl2-mCherry during biotrophic growth of *M. oryzae* at the indicated time points. FYVE localizes as punctate vesicles that colocalize with Pwl2-mCherry in the cytosol. Intensity line graphs of the marked region in the enlarged image of the inset is shown to the right. (C) Single time point images and the sequential images of the corresponding time lapse analysis obtained at 20 s interval. White asterisks indicate the entry of GFP-FYVE positive structure into Pwl2-mCherry marked biotrophy interface. (D) the effect of PI3P kinase inhibitor on the localization of FYVE and invasive hyphal growth. The vesicular and membranous localization of FYVE gets disturbed and is retained in the lumen of enlarged vacuoles; and the BIC formation is abnormal. Pwl2-mCherry is either mislocalized, distorted or disbursed, or appears to split when compared to the solvent/mock control. The effect on the growth of invasive hyphae or BIC formation varies depending on time of addition of the inhibitor. Scale bar = 10 μm.

### MVB-mediated late endocytic trafficking to the biotrophy interface is unconventional

In *M. oryzae*, the proposed unconventional secretion of cytoplasmic effectors was mostly attributed to the lack of Brefeldin A-based pharmacological intervention wherein the biotrophy interface remains unperturbed (Giraldo et al. 2013). Our aforementioned LY294002 treatment to inhibit the PI3-Kinase activity revealed that the intra luminal vesicle formation and MVB are crucial for the focal biotrophy interface formation and proper secretion. Henceforth, along with LY294002, that blocks ILV formation and MVB biogenesis therein, we used Bafilomycin A1 (Baf A1), a specific vacuolar ATPase inhibitor preventing the late stage endolysosomes and analysed their effect on Rab7 dynamics and BIC formation. As expected, treatment with LY294002 significantly reduced the late endosomal structures in the invasive hyphae resulting in larger fused vacuolar structures and aberrant BIC structures. Treatment with Baf A1 though reduced the number of Rab7 vesicles, had no effect on the ILVs and the multivesicular endosomes during the biotrophy and invasive hyphal growth was normal with a proper focal BIC (Figure 5A). The results suggest that the secretion of cytoplasmic effectors to the interfacial zone and the formation of the focal plane for the BIC follows the unconventional vesicle traffic mediated largely through endosomes or the MVBs. To gain additional insight, we assessed the effect of these inhibitors on the localization of the apoplastic effector Bas4 that is known to be affected by BFA. Upon BFA treatment, Bas4-GFP failed to associate with the EIHM and was internalized into the cytosol; whereas Pwl2-mCherry remained unaffected. With Baf A1 and Concanamycin A (Con A) that share the function in inhibiting endosomal and vacuolar fusion, neither the Pwl2 nor Bas4 secretion was affected. Nevertheless, the effect of LY294002 on BIC formation was consistent while it had no effect on Bas4 localization (**Figure 5B**). We conclude that the wMVB adjacent to the BIC of the blast pathogen is involved in endosomal/vesicular trafficking from the invasive hyphae to the biotrophy interfacial complex and it is likely unconventional since it remains unperturbed by BFA treatment.

**Figure 5.**
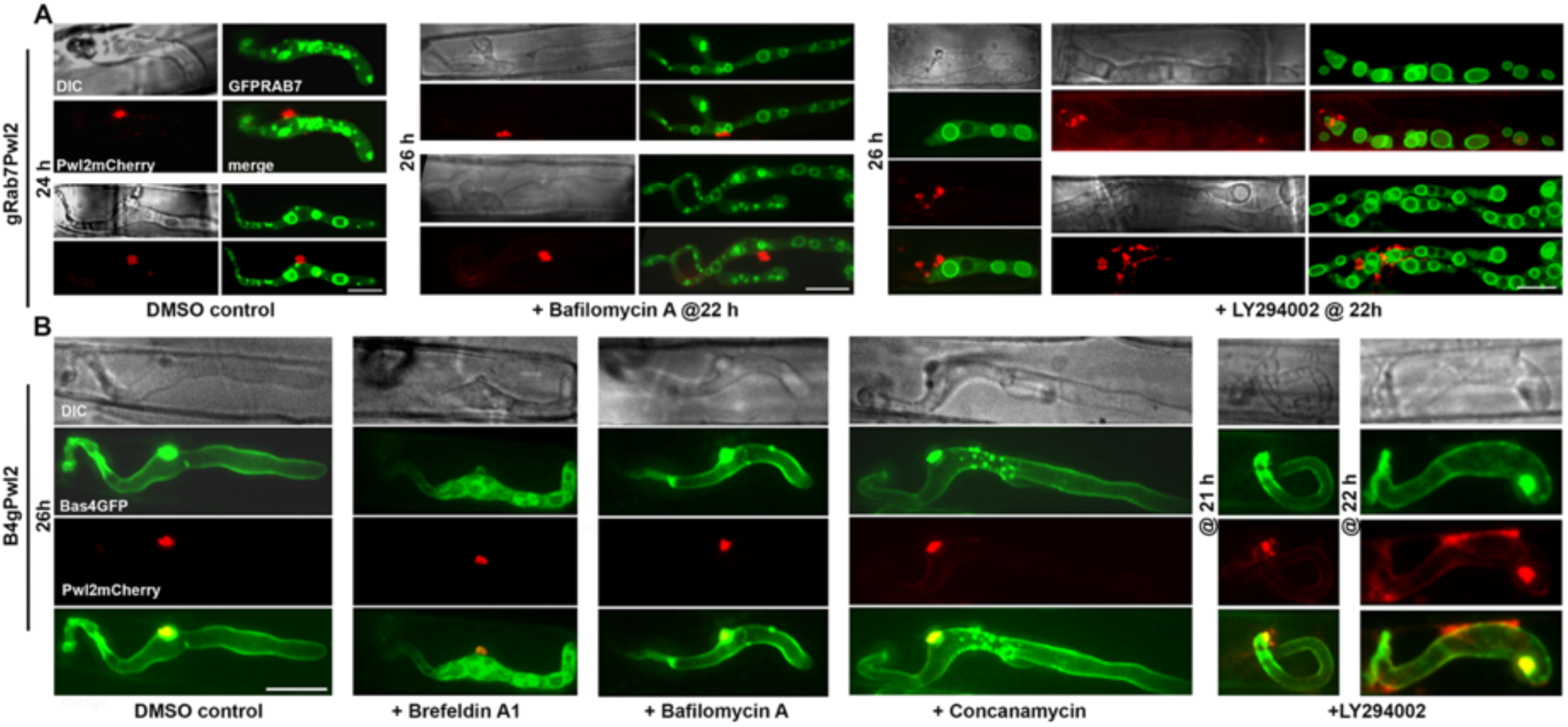
Multivesicular endosomes-mediated late endocytic trafficking to the BIC is non-conventional. (A) Effect of Bafilomycin A and LY294002 on the Rab7 vesicular dynamics and BIC formation in the *gRab7Pwl2* strain. BIC remains unaltered when treated with Baf A, whereas its integrity and proper focal position is perturbed upon inhibition of the PI3P kinase via LY294002. MVB formation is compromised upon treatment with LY294002 leading to significant enrichment of large vacuolar structures that lack intralumenal vesicles. (B) BIC is unaltered when the *Bas4gPwl2* strain is treated with different inhibitors except with LY294002 while Bas4 localization is affected when the conventional ER-to-Golgi trafficking is disrupted with BFA.

### Vta1 marks the MVE membranes and is trafficked to the BIC

As the endosomal vesicles from the MVE form the basis of the unconventional secretion of cytoplasmic effectors in *M. oryzae*, we looked for a protein that could serve as a marker for MVE membrane since Rab7 or FYVE is present in multiple endosomal compartments. In MVB sorting, the reversible membrane association of ESCRT-III is catalysed by Vps4 ATPase activity stimulated by Vta1 that further interacts with other proteins acting during the late stages of the MVB pathway. It is known that upon ESCRT-III disassociation the cargo proteins are found on the membrane invaginations of the MVB and the Vps4/Vta1 complex on the endosomal membrane (**Figure 6A**, Lottridge et al., 2006). We considered Vta1 as an ideal marker for MVB membrane and created the Vta1-mScarlet and Vta1-GFP fusions, since the phenotype associated with Vta1 were not severe, and Vps4 retains the ATPase activity in the absence of Vta1 (Lottridge et al., 2006). Vta1-mScarlet localized to cytoplasmic punctae in the conidia and appressorium, traversing through the germ tube, and was also present on the vacuolar membrane (**Figure 6B, 6C**). Vta1 punctae did not enter the vacuole lumen when stained with CMAC and were present along the membranes, thus implying that they are not destined for degradation therein (**Figure 6C, Video S12**). Vta1-mScarlet colocalized with GFP-Rab7 vesicles at the membranes of the Rab7-marked late endosomes, specifically the MVBs with the highly dynamic intraluminal vesicles in gRab7Vta1mSc (**Figure 6D, 6E**). Further, time lapse analysis and surface rendered imaging of Vta1gPwl2mC strain clearly showed that Vta1-GFP punctae were abundant near the BIC, colocalized with Pwl2-mCherry (**Figure 7A, 7B**) and Vta1 is continuously trafficked to the BIC (**Videos S13-S15**). However, such association of Vta1 was not evident with Bas4-mCherry that marks the EIHM, thus confirming the specificity of Vta1 localization to the cytoplasmic effectors (Figure S7A).

**Figure 6.**
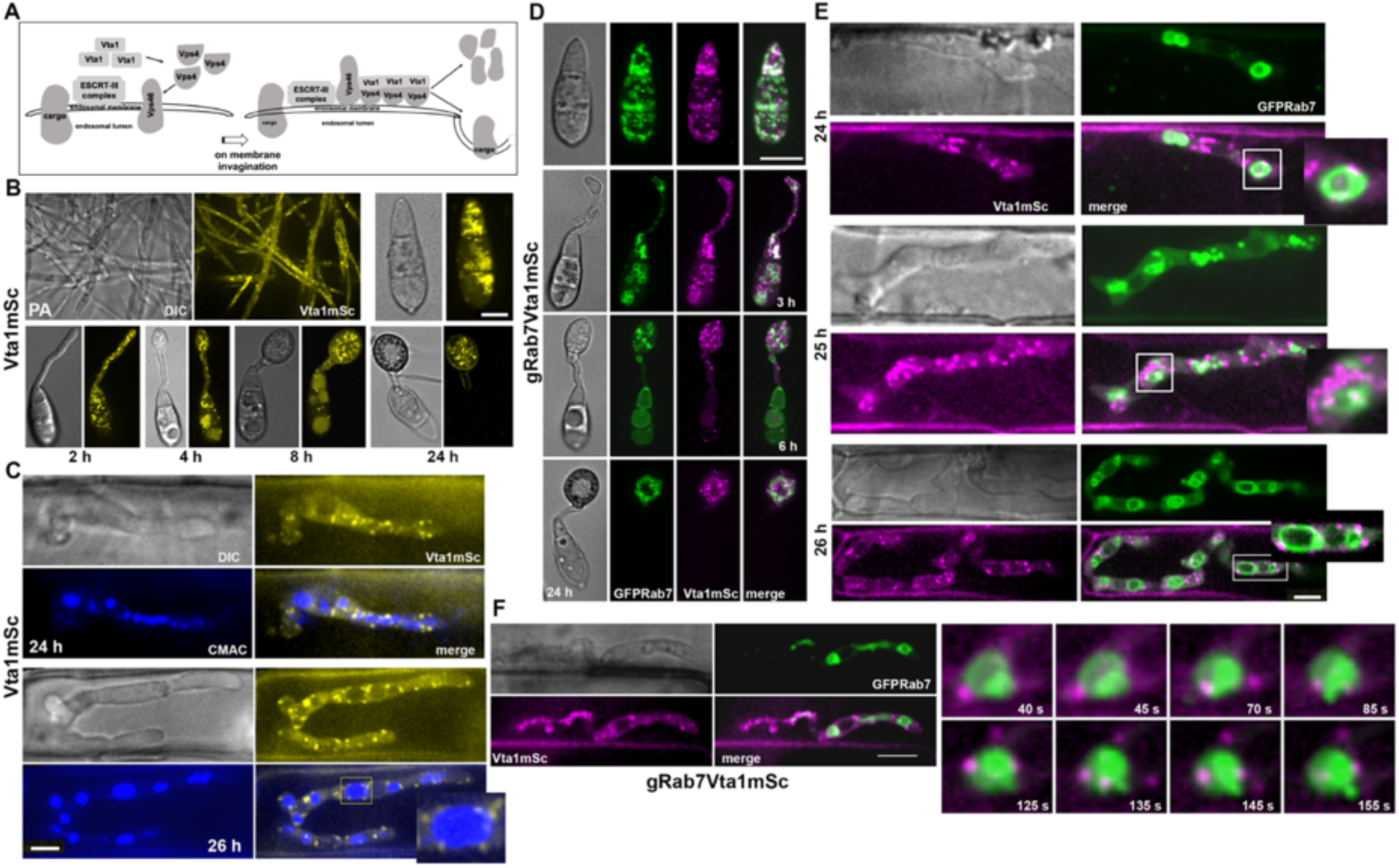
Vta1 marks the multivesicular endososomal or MVB membranes. (A) Schematic rendition of the shift in localization of Vps4 and its cofactor Vta1 associated with the function of the ESCRT-III complex and membrane invagination (modified from Lottridge et al., 2006). (B) Localization of Vta1-mScarlet in the vegetative mycelia and in the infection structures at different time points in *M. oryzae*. Vta1 localizes to vesicular punctae in the cytoplasm. (C) Co-localization of Vta1mSC with the vacuoles. Vta1 localizes as punctate vesicles on the vacuolar membranes whose lumen is marked with CMAC. (D) Vta1-mScarlet co-localizes with GFP-Rab7 late endosomes during conidiation and appressorium formation as observed at different time points in *gRab7Vta1mSc* strain. (E) Vta1 localizes to MVB membranes marked by GFP-Rab7 in the invasive hyphae of strain. The insets depict enlarged images of the sections outlined in white, and clearly show Vta1 and Rab7 colocalization on MVBs. (F) Images showing the localization of Vta1-mScarlet and GFP-Rab7 in the invasive hyphae. Selected time points of the time lapse obtained at 5 sec interval showing the mobility of Vta1-mScarlet along the multivesicular endosomal membranes marked by GFP-Rab7. Scale Bars represent 10 um.

**Figure 7.**
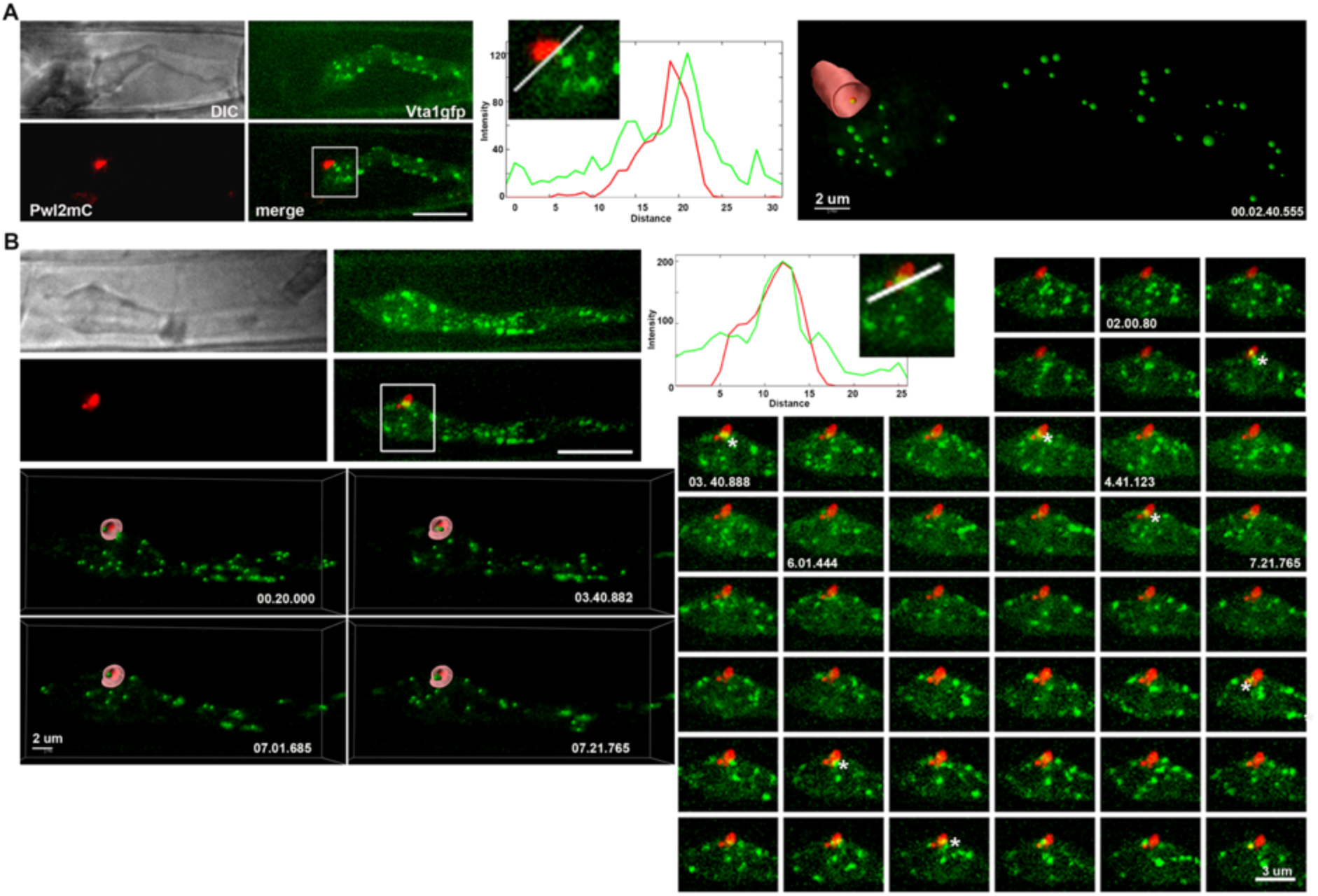
Vta1 resident MVB vesicles are trafficked to the BIC. (A) Vta1-GFP punctae are abundant near the biotrophy interface marked with Pwl2-mCherry. The enlarged images of the section (white rectangle) and line graphs representing the intensities along the marked (white line) region are shown. The surface rendered 3D image on the extreme right depicts Vta1 punctae entering the BIC. (B) Surface rendered 3D images, and the sequential micrographs obtained in a time lapse at 20 sec intervals (z=13 at 0.3 u distance) are shown for *Vta1gPwl2mC* strain at 26 h. The images clearly show Vta1-GFP entering the Pwl2-mCherry marked biotrophic interface (white asterisks). Scale bars represent 10 μm unless indicated otherwise.

Furthermore, Vta1 deletion mutant did not show any phenotypic or pathogenesis defects in *M. oryzae* (**Figure S7B**) with minimal or negligible effect on the ILV formation of MVBs and the BIC biogenesis being comparable to the WT (**Figures S7C, S7D**). Live cell imaging with Vta1 revealed that the unconventional vesicular trafficking involved in cytoplasmic effector secretion to the BIC and hence proper focal BIC formation is mediated through the MVBs. Based on the results presented here, we propose a model for establishment of proper biotrophy interface and the associated secretion machinery for effectors in which the *M. oryzae* MVBs of the late endocytic and endolysosomal pathway are redirected to or near the BIC, fuse with the plasma membrane and release their intralumenal content with phosphoinositides into the BIC as exosomes (**Figure 8**) from where it may be internalized by the living host cells via clathrin-mediated endocytosis (Garcia et al., 2023) leading to successful onset of the devastating blast disease in rice.

**Figure 8.**
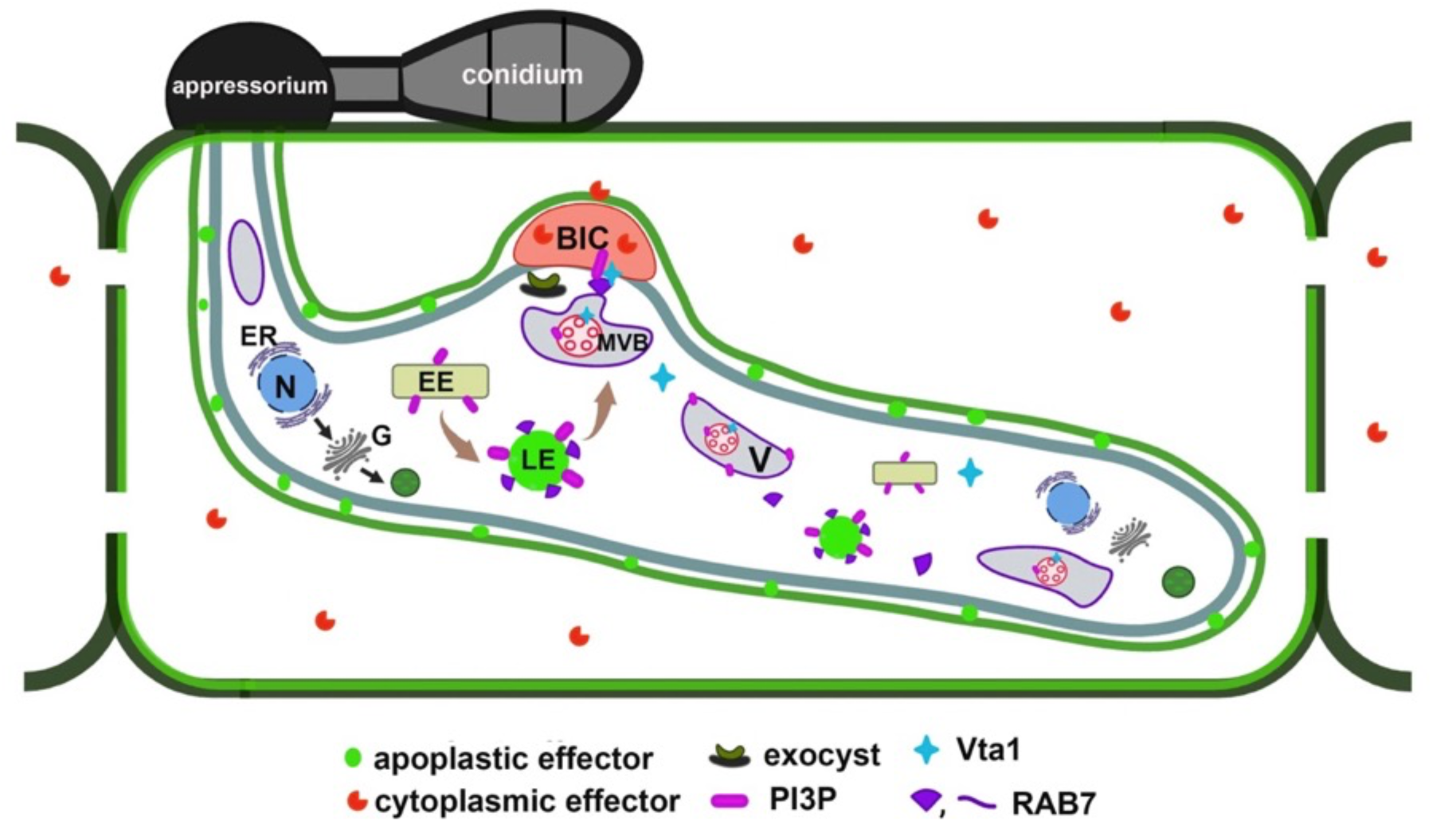
Proposed model for multivesicular bodies dependent non-conventional secretion of cytoplasmic effectors in *M. oryzae*. Rab7, the late endosomal (LE) protein marking the late endocytic machinery matures into multivesicular bodies (MVBs) with intralumenal vesicles (ILV) rich in PI3P that fuse with the BIC membrane and releases their ILV contents in the form of extracellular vesicles (EVs) into the biotrophy interface. Perturbation of this vesicular membrane trafficking system disrupts the BIC formation. Vta1, the cofactor of the vacuolar ATPase Vps4 marks the membrane of MVB and is also trafficked to the BIC. The vesicles secreted to the BIC are composed of endocytic components Rab7, Vta1 and phospholipid PI3P in the fungal pathogen. The model proposes the involvement of such pathogen endocytic machinery in the proper formation and maintenance of the interfacial zone BIC and its role in spatiotemporal regulation and the timely secretion of the cytoplasmic effectors. EE-early endosomes, LE – late endosomes, MVB – multivesicular bodies, V-vacuole, N-nucleus, ER – endoplasmic reticulum, G – Golgi body.

## Discussion

We show that the Rab7 GTPase marks the late endocytic and vacuolar compartments in *M. oryzae,* thus denoting the unique endomembrane systems the MVBs or late endosomes with the intralumenal vesicles. We used live cell imaging of fluorescently labelled *M. oryzae* proteins and effectors to dissect such endomembrane trafficking at and to the host interface, and show that late endocytic trafficking, which normally drives cargo proteins from the cell surface to the vacuole, is diverted/rerouted towards the biotrophy interface during *in planta* growth. The EIHM marked by the apoplastic effectors at any stage does not contain Rab7 whereas the biotrophy interfacial complex does. The Rab7 vesicles traffic to the BIC, thus indicating the importance of Rab7 in cytoplasmic effector secretion in the *Magnaporthe*-Rice interaction zone. We further show that this rerouting is important for focal BIC formation and consequently the pathogenicity of *M. oryzae.* The Rab7 dominant negative variant was incapable of MVB formation and remained cytoplasmic, thereby showing inefficient cytoplasmic effector secretion that results in reduced pathogenicity. This implies that proper focal BIC formation and the translocation of cytoplasmic effector Pwl2 thereto is important for the pathogenicity of *M. oryzae* (Figure 2). Both mutant analysis and live imaging results indicated that Rab7 plays a crucial role in proper invasive hyphal development as well as cytoplasmic effector secretion. However, the biotrophy interface *per se* does not contain any vacuolar identity as indicated by the failure to be stained with the CMAC dye. However, it is the ILV that gets trafficked to the BIC or host interface (Figure 3). We further confirmed this observation using FYVE to label vesicles enriched in PI3P, notably the endosomes and MVBs of the late endocytic pathway (Robinson et al., 2008; Gao et al., 2014).

A growing body of evidence indicates an important role of phosphoinositides in many aspects of microbial colonization of hosts and in particular PI3P-mediated effector entry is widespread in fungal pathogens of plant, animal and humans (Kale et al., 2010; Bhattacharjee et al., 2012; Hu et al, 2008; Plett et al., 2011). The pathogen produced PI3P aids in infection such as to promote host entry of RXLR effectors and is required for the full virulence of *Phytophthora infestans* (Liu et al., 2013). Blocking PI3P binding inhibited effector entry and significantly diminished the virulence; and thus PI3P has been demonstrated as an excellent disease control target for these pathogens ( Liu et al., 2013, Zhou et al., 2021). In addition, recent studies have uncovered a differential and specific recruitment of PI4P, and PI(4,5)P2 though not PI3P, to the host-derived membranes at the biotrophic interface (Shimada et al., 2019, Qin et a., 2020, Li Qin and Wei 2022) that likely signal the onset of the defense response. We have shown here that treatment of conidia with LY294002 that inhibits PI3P biosynthesis results in abnormal BIC formation and significantly reduces the virulence (Figure 4). Nevertheless, it is possible that *M. oryzae* deploys the cytoplasmic effectors to boost the late endocytic pathways, helps recruit endomembranes for biotrophy interface biogenesis to establish a conduit for countering the host immunity via secreted effectors or metabolites, as a potential pathogen strategy to promote infection (Bozkurt et al., 2015, Rausche et al., 2020, Peter et al., 2021).

The presence of PI3P FYVE vesicles at the interfacial complex suggests that the late endocytic / endolysosomal trafficking diverted and rerouted to the BIC might involve MVBs and the exosomes like EVs that form a distinct unconventional secretion pathway for virulence factors by such fungal pathogens. The cytoplasmic effectors in *M. oryzae* are secreted via an non-conventional route, and the BIC and hence the cytoplasmic effectors remain unperturbed when the conventional ER-to-Golgi secretion is blocked using Brefeldin A (Giraldo et al., 2013). We further extended this pharmacological approach to other inhibitors and show that *M. oryzae* employs the MVB routed extracellular vesicles for the proper biogenesis and/or establishment of the interfacial complex and to secrete the cytoplasmic effectors. Inhibitor-based disruption of the conventional ER-Golgi trafficking or the conventional trafficking of vesicles to the vacuole for degradation or autophagy resulted in normal BIC formation and pathogenicity of *M. oryzae*, even though there were changes in the normal vesicular structures formed by Rab7 (Figure 5). However, when the formation of multivesicular endosomes is blocked by LY294002, it affected the BIC formation and resulted in reduced hyphal proliferation even though the secretion of apoplastic effector Bas4 remained normal (Figure 5). These results ruled out the involvement of conventional ER-Golgi trafficking or autophagy in the effector secretion process and illustrated that the multi-vesicular bodies or extracellular vesicles are involved in the unconventional secretion of cytoplasmic effectors.

Vesicular secretion is an important mechanism in fungi to deliver intracellular molecules to extracellular space, in particular factors related to virulence and fungal EVs are a mechanism for non-conventional secretion, that have potential functions in host-pathogen interactions (Albuquerque et al., 2008). Studies of fungal and oomycete biotrophic interfaces with abundant MVBs in pathogen and host cytoplasm have revealed such conduits as sites of active secretion involving both partners across the interfacial zone (Yin and Valent 2013; Micali et al, 2011; Baur et al., 2004; Mims et al., 2004) though the nature and composition of the interfaces varies significantly. Recently, in *C. higginsianum*, another hemibiotroph that shares a similar infection strategy as *M. oryzae,* membrane-bound vesicular structures were observed in the paramural space between the fungal cell wall and plasma membrane of biotrophic hyphae, such that EVs could be easily extracted from the fungal spheroplasts (Rutters et al., 2022). These studies used electron microscopy and proteomics-based characterization to investigate the structural integrity and constituents of the extracted EVs. However, based on TEM evidence alone and in the absence of specific molecular markers, we are unable to determine whether the vesicles observed trult represent exosomes derived from the fusion of fungal MVBs or whether these originate from host vesicles. In our study, apart from using Rab7 or FYVE probe as cellular markers, we describe here Vta1 as a specific marker for MVBs and show that late endolysosomal or multi-vesicular trafficking is involved in the secretion of cytoplasmic effectors in rice blast. Vta1 is a crucial component of MVB machinery regulating the function of the AAA-ATPase Vps4, that catalyses the disassembly and dissociation of ESCRT III complex from the endosomal membranes. Since ATP hydrolysis is irreversible, the regulation of Vps4 ATPase activity is critical for proper functioning of the MVB sorting reaction and machinery. Vta1 punctae that partially colocalize with Rab7 on the membrane counterparts were also evident on the membranes of intralumenal vesicles, whereas such punctate structures were absent at the cytoplasmic face of the ILVs. Furthermore, such Vta1 punctae were abundant in the cytoplasm near the biotrophic interfacial complex or zone, and Vta1 was found to be trafficked to the BIC periodically therefrom (Figures 6 - 7).

Although abundant in the cytoplasm, Vta1 was never found to associate with the extra-invasive hyphal membrane marked by Bas4, and hence we concluded Vta1 to be exclusively specific to secretion of cytoplasmic effectors only. Interestingly, the invasive hyphae formed by the Vta1 mutant are not impaired thus indicating a potential redundancy of the protein function. Alternately, it is also likely that Vps4 exhibits ATPase activity in the absence of Vta1 and thus the phenotypes associated with the loss of Vta1 may not be severe as those found in the *VPS4* deletion mutant in yeast (Yeo et al, 2003; Shiflett et al., 2004). Taken together, we present sufficient evidence for an MVB-mediated exosome or extracellular vesicle trafficking for the unconventional secretion of the cytoplasmic effectors to the biotrophy interface in rice blast (Figure 8) wherefrom such proteins may be internalized into the living rice cells through clathrin-mediated endocytosis (Garcia et al., 2021). The possible existence of such an alternate vesicle transport route, distinct from the canonical ER/Golgi-based pathway, for the secretion of cytoplasmic effectors is pertinent for *M. oryzae* (Khang et al., 2010; Giraldo et al., 2013) since the rice blast cytoplasmic effectors lack a canonical secretion signal as predicted for the pathogenic oomycete counterparts.

In summary, our results provide evidence for an active Rab7 and endolysosomal vesicle trafficking involving PI3Ps and MVBs that is unconventional, is critical for focal host-pathogen interface formation, and most importantly for establishing the biotrophic phase in the infection cycle of *M. oryzae*. Unravelling the mechanism of cross-kingdom cytoplasmic effector secretion by a distinct pathway is momentous for blast research since the interfacial zone is important not only in effector delivery but also for specific sequestration therefrom by the host. Future studies will help further elucidate the mechanisms and involvement of the multi-vesicular late endosomes and endolysosomal compartments in trans-kingdom effector delivery. Likewise, the manipulation of host endomembrane systems will prove pivotal in controlling rice blast outbreaks and also in gaining insights into the evolution/adaptation of exosomes or extracellular vesicles in membrane trafficking associated with infection-related secretion in fungal pathogens to control the resultant devastating diseases in important crops such as rice and wheat.

## Materials and Methods

### Strains, plasmids and cultural conditions

*Magnaporthe oryzae* B157 was used as the wild-type isolate in this study. The fungal strains were maintained on Prune juice agar (PA) or Complete medium (CM) for 7-10 d at 28°C and stored as dry filter paper stocks at −20°C. All plasmids were constructed by standard molecular cloning procedures and confirmed by restriction enzyme digestions and nucleotide sequencing. Detailed cloning information including the list of plasmids, DNA manipulation and transformation of *M. oryzae* strains using standard *Agrobacterium tumefaciens* mediated transformation is provided in the Supplementary material (Table S1, S2 and S3). Fungal mycelial blocks were inoculated in CM medium and cultured for 2-4 days for DNA extraction and the transformants were confirmed by locus-specific PCR and sequencing. In all cases, at least two strains were characterised and imaged for confirmation and the results from either or both the strains are presented here.

### Conidial germination, appressorium formation and plant infection assays

For conidia production, strains were cultured on PA media for 2 days at 28°C in the dark and then transferred to light and grown for 5 to 7 days. The conidia were harvested on the 7^th^ day and inoculated on coverslips or cover glass dish for live cell confocal imaging. For infection assays, spore suspension (1×10^5^ conidia/ml) of the indicated strain was inoculated on either barley (8 day old) or rice leaves (variety CO39, 14 to 21 day old) and incubated at 25°C on a 16h day / 8h night cycle for 5 days in a growth chamber. For live imaging of invasive growth spores were inoculated on rice leaf sheath maintained on 1% agar medium incubated at room temperature under humid conditions for the required period and observed using laser scanning confocal microscopy. To observe the infection at different time points, barley leaves inoculated with spore suspension were cut into pieces at the required time points and decolorized with methanol before visualizing under bright field microscope.

### Chemical inhibitors and microscopy

For treatment with inhibitors, the spore suspension was inoculated to leaf sheaths and allow infection for 22h after which the whole inoculated tissues were treated with the inhibitors. Stocks of Brefeldin A, Concanamycin, Bafilomycin A1 and LY294002 were prepared in DMSO and diluted to required concentrations in 1% DMSO solution in which the sheath samples were incubated for mentioned time points. To observe the cytological changes during biotrophy, the vacuolar lumen was stained with CMAC (7-amino-4-chloromethylcoumarin, Cat. #C2110, Invitrogen) used at a final concentration of 10 μM in DMSO. Bright field microscopy was performed with an Olympus BX51 microscope (Olympus, Tokyo, Japan) using a Plan APO 100X/1.45 or UPlan FLN 60X/1.25 objective lens. Images were captured with Photometrics CoolSNAP HQ camera (Tucson, AZ, United States) and processed using MetaVue (Universal Imaging, Downingtown, PA, United States) software associated with the system. Live cell imaging was performed in a confocal microscope equipped with a Yokogawa CUS-X1 spinning-disk and a 100 x/1.4 NA oil lens objective. Micrographs were captured in a 16-bit Orca-Flash 4.0 scientific complementary metal oxide semiconductor (sCMOS) camera (Hamamatsu Photonics KK). The excitation/emission wavelengths for GFP, mCherry and mScarlet and CMAC were 488 / 500 −550 nm, 561 / 570-620 nm and 405/ 425-475 nm respectively. Image processing, time lapse analysis and surface rendering were performed using Fiji (https://imagej.net/Fiji) or Imaris (Bitplane) and Adobe Photoshop 21.1.2 (Mountain View, CA, United States).

## Supplementary Data

**Supplementary Tables, Supplementary Figures and Supplementary Movies** are available at: https://doi.org/10.5281/zenodo.14038222 or https://zenodo.org/records/14038222

---

**Table S1.**
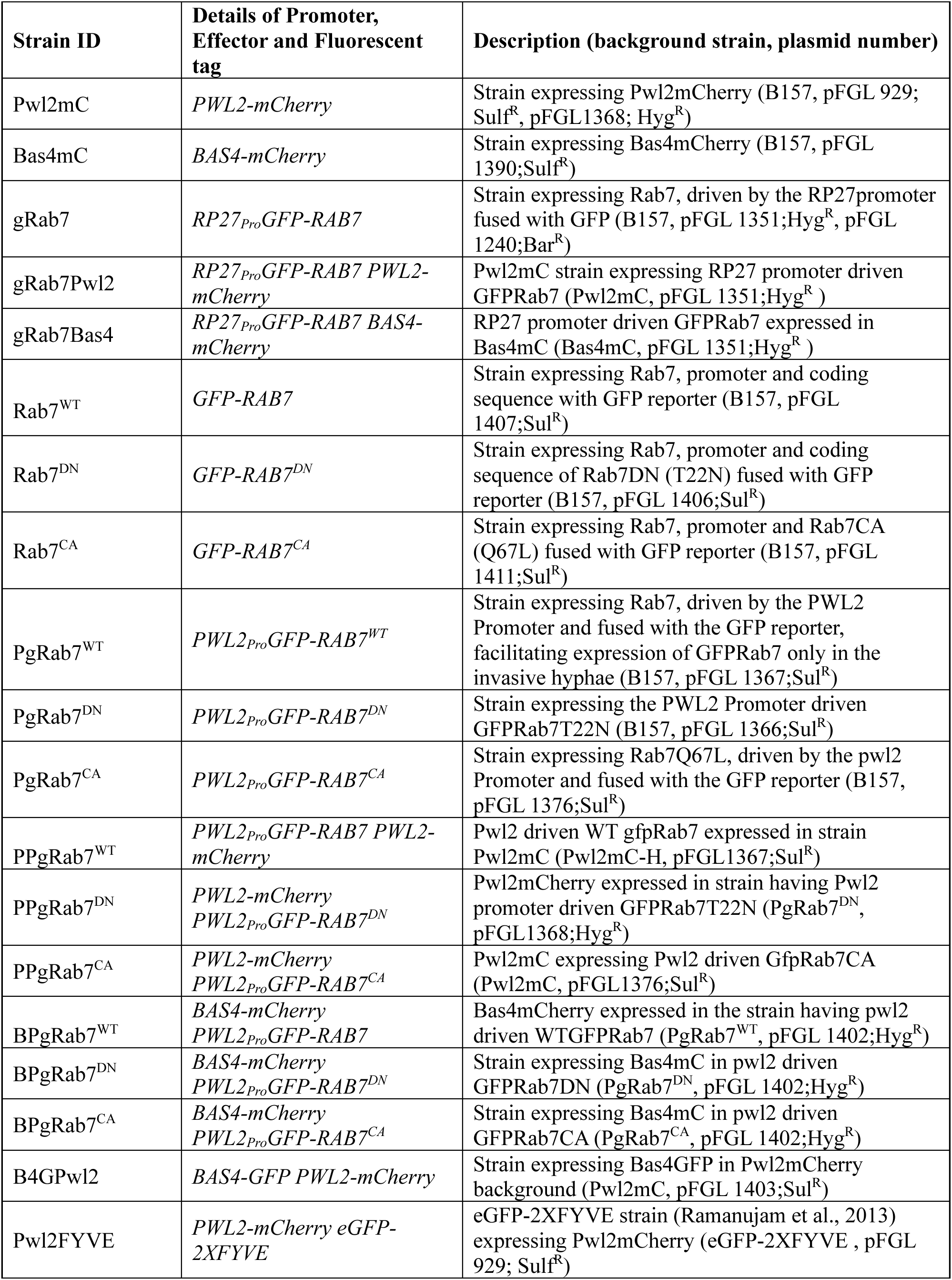
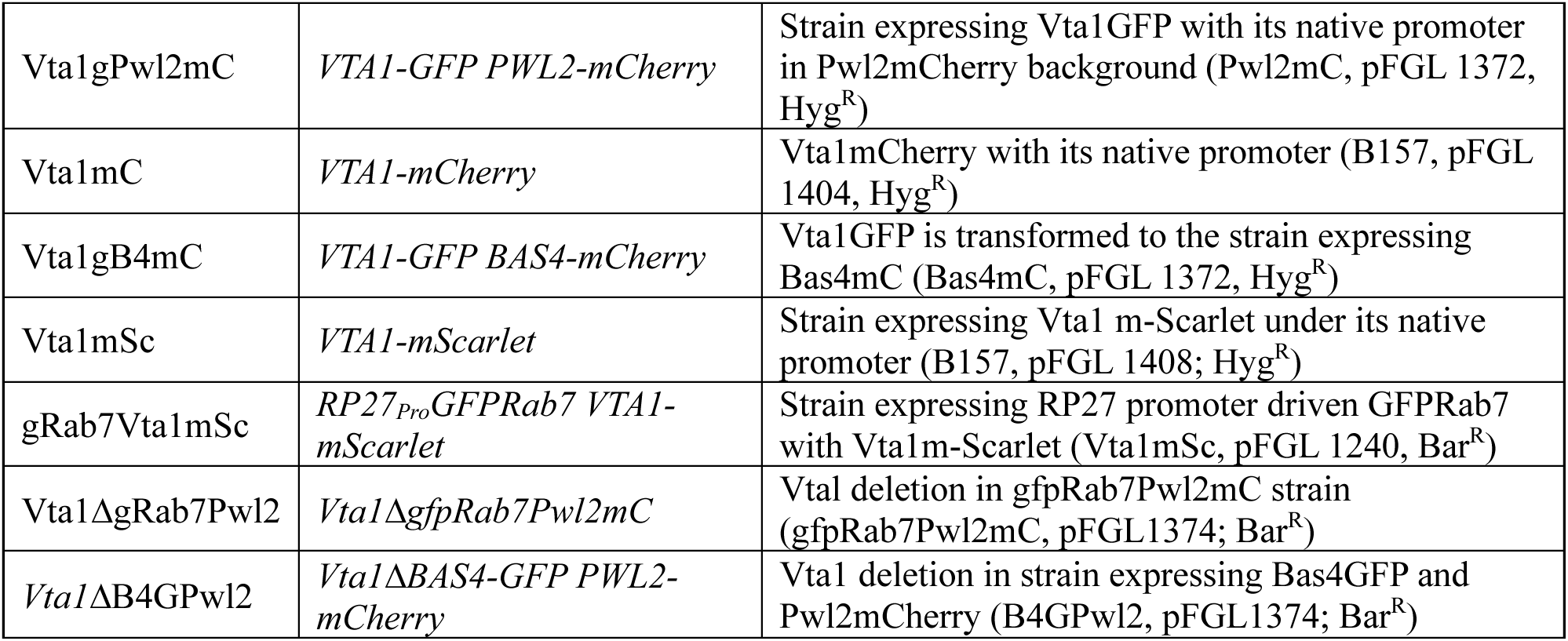
Details of the *M. oryzae* strains generated in this study.

**Table S2.** Key plasmids used for *M. oryzae* transformation in this study.

| Plasmid | Description |
| --- | --- |
| pFGL 929 | Pwl2 promoter and entire coding sequence fused with mCherry reporter gene and the TrpC terminator sequence at its C terminal. The entire 1.3kb of Pwl2 gene, PCR amplified with EcoRI/ KpnI restriction sites and mCherry with terminator obtained through kpnI/XbaI digestion of pFGL810 were digested and sequentially cloned into the respective sites of pFGL 920 (EcoRI/XbaI) with sulfonyl urea resistance marker (Sul<sup>R</sup>) |
| pFGL1368 | Pwl2mCherry – Pwl2promoter and entire coding sequence with the translational fusion of mCherry reporter gene obtained as EcoRI/BamHI fragment from pFGL929 was cloned into the respective sites of pFGL972, the plasmid containing Trpc terminator and hygromycin resistance marker (Hyg<sup>R</sup>) |
| pFGL 1390 | Bas4 promoter and entire coding sequence fused with mCherry reporter gene and the TrpC terminator sequence at its C terminal. The fragments Bas4promoter + ORF (EcoRI/KpnI), mCherry with terminator (KpnI/XbaI) were cloned into the respective sites (EcoRI/XbaI) of pFGL1010 that contains the iLV locus for sulfonyl urea resistance (Sul<sup>R</sup>) |
| pFGL 1240 | Rab7 entire coding sequence with eGFP reporter gene fused at its N terminal driven by the constitutive RP27 promoter in pFGL97 containing Glufosinate ammonia resistance (Bar<sup>R</sup>) – plasmid published in Ramanujam et al., 2011. |
| pFGL 1351 | The plasmid pFGL1240 was replaced with Hygromycin resistance from pFGL44 as BamHI/PstI fragment (Hyg<sup>R</sup>) |
| pFGL 1407 | Rab7 promoter (EcoRI/KpnI) and entire coding sequence fused with GFP (PstI) obtained as PCR products are digested with the respective enzymes and subsequently cloned into pFGL820 containing sulfonyl urea resistance (Sul<sup>R</sup>) |
| pFGL 1406 | Rab7 promoter (EcoRI/KpnI) and entire coding sequence with single nucleotide substitution at T22N (PstI), the dominant negative version (DN-GTPase dead)) fused with GFP reporter at its N terminal. The fragments are obtained as PCR products were digested with and subsequently cloned into the respective enzyme sites of pFGL820 containing sulfonyl urea resistance (Sul<sup>R</sup>) |
| pFGL 1411 | Rab7 promoter and entire coding sequence with single nucleotide substitution at Q67L, the constitutive active mutant version (CA-GTP bound state) fused with GFP reporter at its N terminal. The PCR fragment Rab7Q67L (PstI) is replaced for Rab7 in pFGL1407 containing sulfonyl urea resistance (Sul<sup>R</sup>) |
| pFGL 1367 | Rab7 the entire coding sequence fused with the GFP reporter at its N terminal driven by the pW12 Promoter that facilitate expression of GFP Rab7 during the invasive hyphal stage alone. The full fragment is cloned by sequential enzyme digestion and cloning into respective sites (EcoRI/XbaI) of pFGL1010 containing the ILV locus of sulfonyl urea resistance (Sul<sup>R</sup>) |
| pFGL 1366 | Rab7T22N coding sequence fused with GFP at its N terminal driven by pwl2 promoter. Constructed as pFGL1367 but containing Rab7DN coding sequence (Sul<sup>R</sup>) |
| pFGL 1376 | Rab7Q67L coding sequence fused with GFP at its N terminal driven by pwl2 promoter. Constructed as pFGL1367 but containing Rab7CA coding sequence in the place of Rab7 (Sul<sup>R</sup>) |
| pFGL 1402 | Bas4 promoter and entire coding sequence fused with mCherry reporter gene and the TrpC terminator sequence at its C terminal. The entire cassette as EcoRI/XbaI fragment from pFGL 1390 was transferred to pFGL44 containing a hygromycin resistance into the respective enzyme sites (Hyg<sup>R</sup>) |
| pFGL 1403 | Bas4 promoter and entire coding sequence fused with GFP reporter gene and the TrpC terminator sequence at its C terminal. Bas4mC clone (pFGL1390) was digested with KpnI/ BamHI and subsequently cloned with GFP digested with KpnI / BamHI (Sul<sup>R</sup>) |
| pFGL 1372 | Vta1 entire coding sequence with an eGFP translational fusion at its C terminal with the Trpc terminator, driven by its native promoter. Vta1ORF (EcoRI/KpnI) and Vta13' UTR (PstI) are cloned into the respective enzyme sites of pFGL972 containing the eGFP and Trpc terminator sequences with Hygromycin resistance (Hyg<sup>R</sup>) |
| pFGL1374 | Vta1 deletion with glufosinate ammonia resistance (Bar<sup>R</sup>). The upstream 5'UTR (EcoRI / XmaI) and downstream 3'UTR (PstI) PCR products of Vta1 is digested and sequentially cloned into the respective sites of pFGL97 with Bar resistance. |
| pFGL 1404 | Vta1 coding sequence with mCherry and Trpc terminator. mCherry replaced the GFP sequence in pFGL1372 with Hygromycin resistance (Hyg<sup>R</sup>) |
| pFGL 1408 | Vta1 coding sequence with m-Scarlet and Trpc terminator at its C terminal. m-Scarlet replaced the GFP sequence in pFGL1372 with Hygromycin resistance (Hyg<sup>R</sup>) |

**Figure S1.**
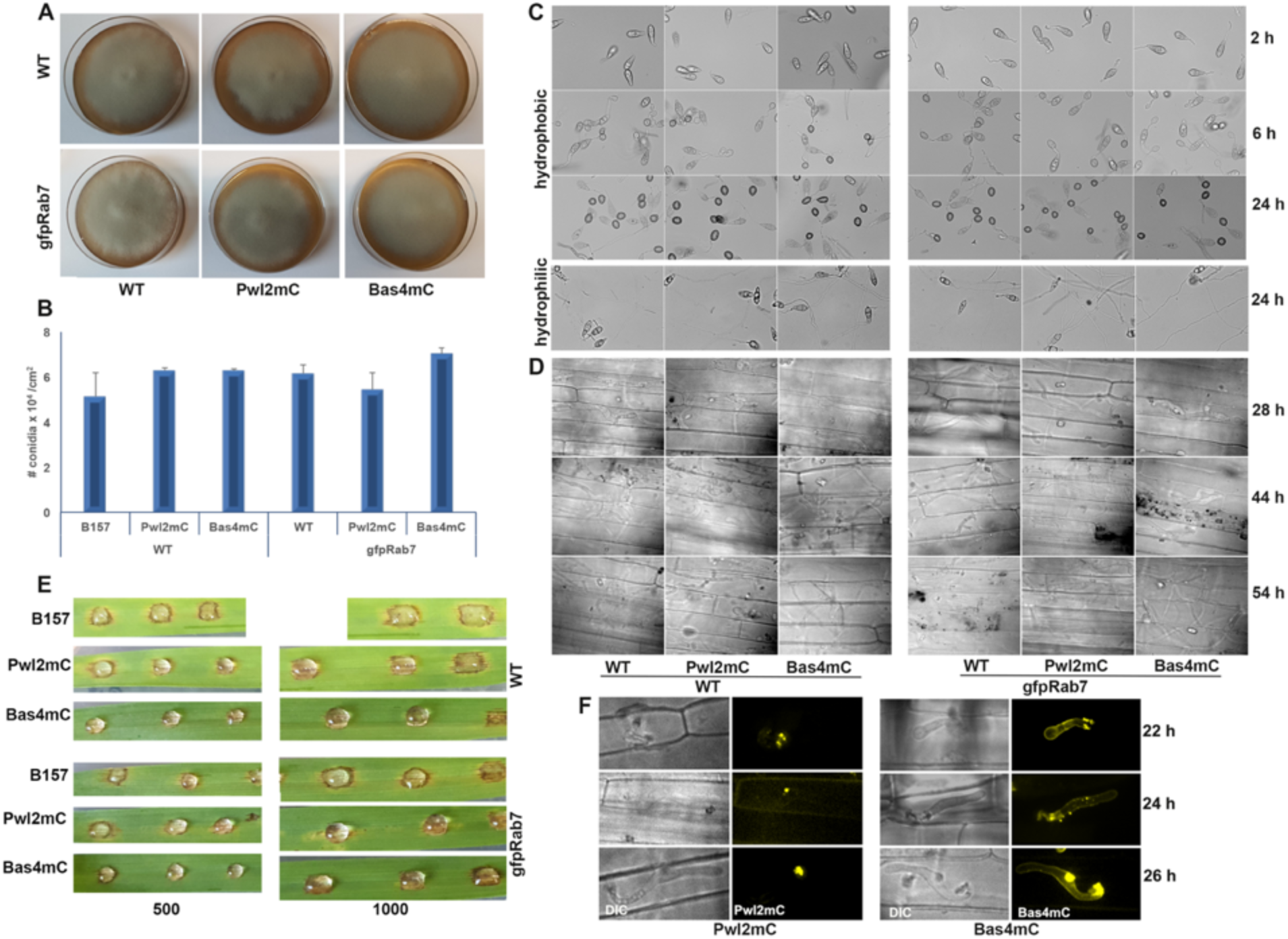
Characterization of strains co-expressing GFP-Rab7 with Pwl2-mCherry (gRabPwl2) or Bas4-mCherry (gRabBas4) representing a cytoplasmic and apoplastic effector, respectively. (A) Colonies showing the growth of GFP-Rab7 strains expressed in the WT B157 background, Pwl2-mCherry and Bas4-mCherry strains compared with the corresponding WT. (B) Bar graph showing the conidiation levels in the indicated strains. Each column represents mean ± standard error (S.E) of three independent replicates and the experiment was repeated twice. (C) Images showing the conidial germination and appressorium formation in the indicated strains under hydrophobic or hydrophilic conditions. (D) Micrographs showing the invasive hyphal growth in rice leaf sheath at indicated timepoints. (E) Blast disease lesions produced by GFP-Rab7 Pwl2-mCherry or Bas4-mCherry strains compared to the corresponding wild type in barley (top) and rice leaves (bottom). (F) Confocal images showing the localization of Pwl2-mCherry and Bas4-mCherry in the respective strain co-expressing GFP-Rab7.

**Figure S2.**
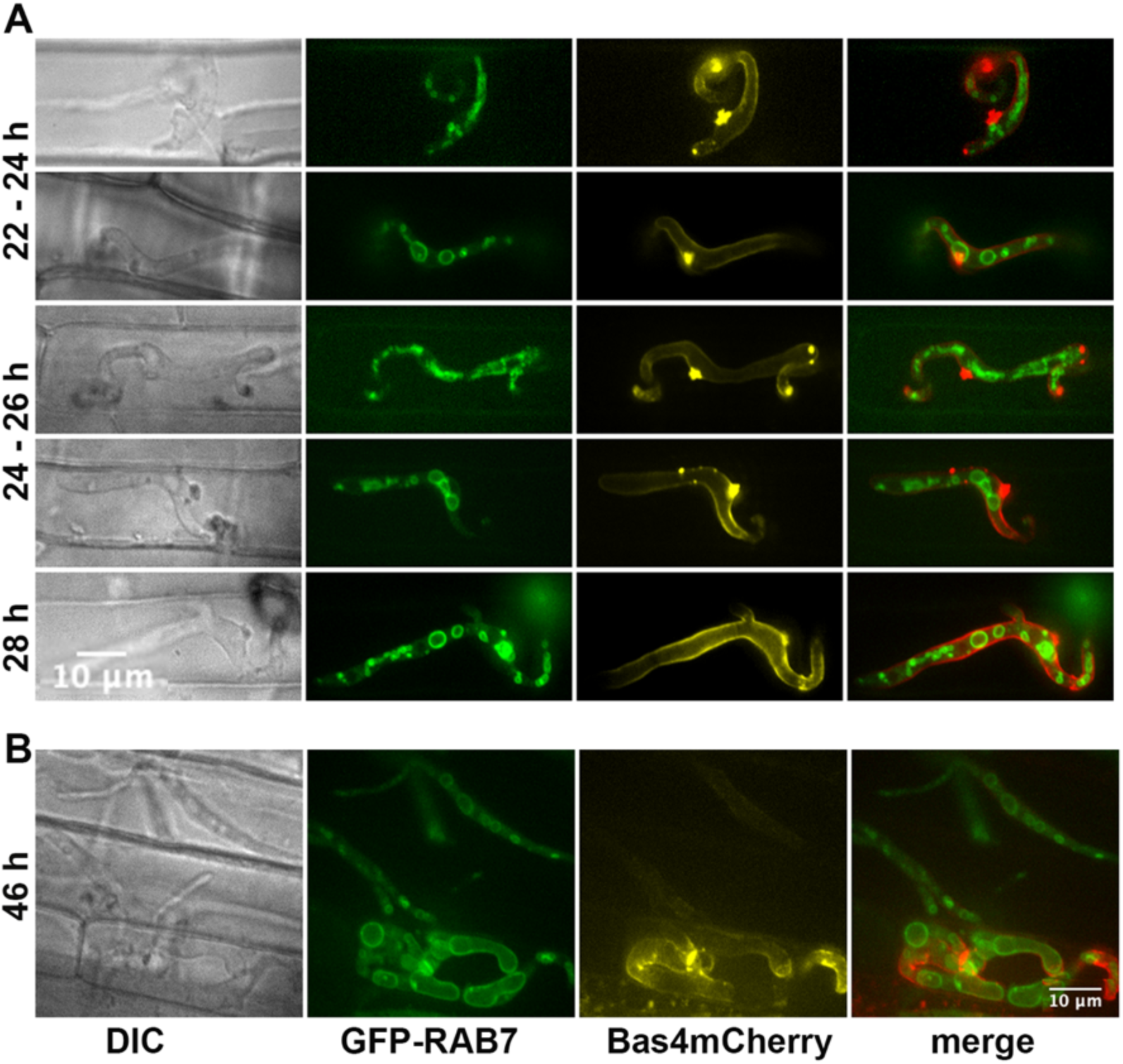
Localization of GFP-Rab7 and Bas4-mCherry in the biotrophic hyphae at (A) earlier time points during the tip, side and focal BIC stages and (B) later time points (46 h) in gRab7Bas4. Bas4-mCherry localizes to the extra-invasive hyphal membrane and GFP-Rab7 localizes to the vesicles and vacuolar membrane but does not co-localize with Bas4-mCherry in the EIHM.

**Figure S3.**
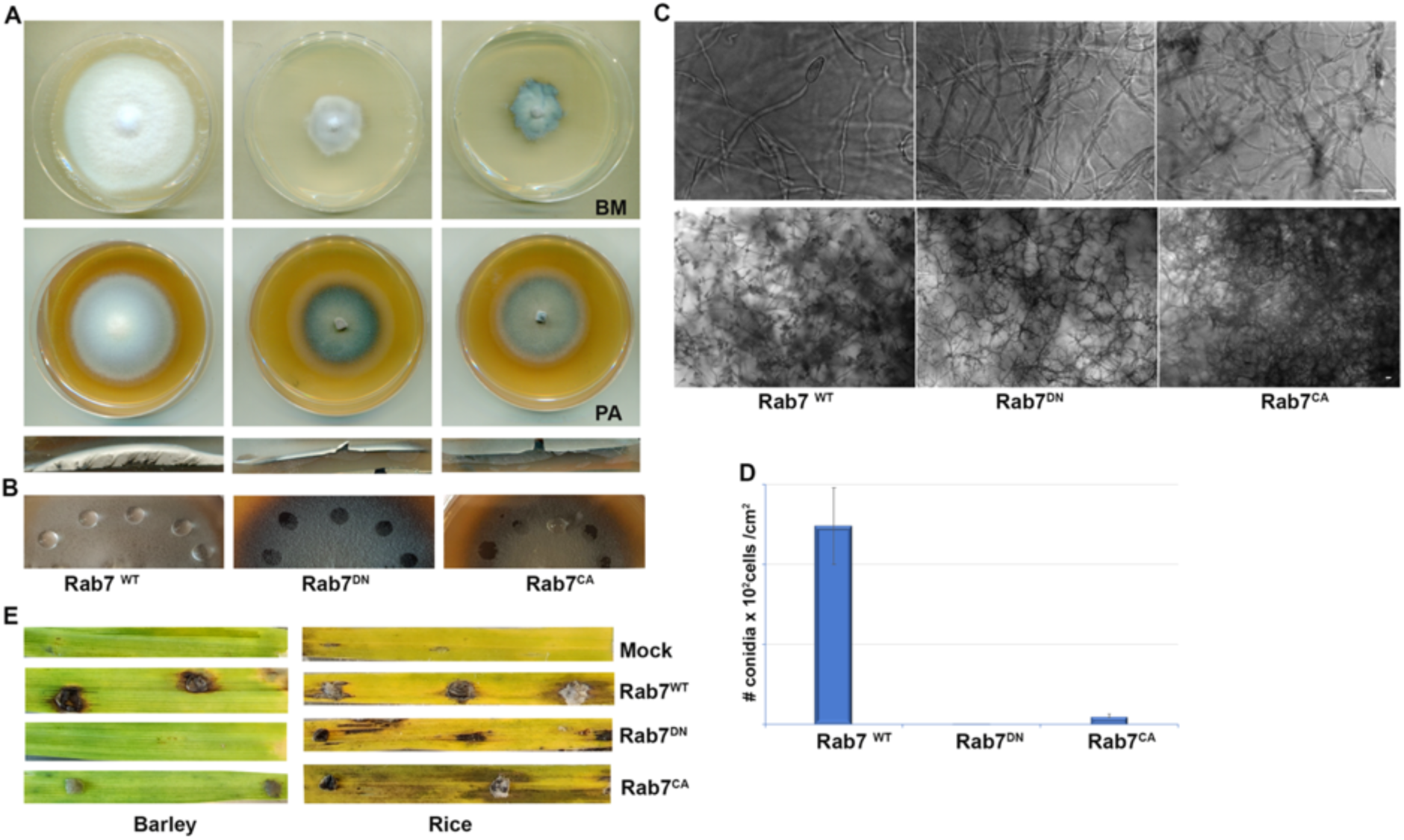
Characterization of Rab7 mutants, the dominant negative (Rab7^DN^) and constitutive active (Rab7^CA^) along with WT (RAB7^WT^). (A) Colonies showing the radial and aerial growth of Rab7^WT^ or Rab7^DN^, or Rab7^CA^ mutant in Basal medium (BM) or Prune juice agar (PA). (B) Surface hydrophobicity of Wild type and Rab7 mutants in PA. (C) Micrographs showing the conidiophore (top) production and conidiation (bottom) in wild type and GFP-Rab7 mutants. (D) Quantification of conidia produced by WT and the indicated Rab7 mutants. Each value is the mean ± S.E of three individual replicates. (E) Pathogenicity of the mycelial plugs of WT and GFP-Rab7 mutants in barley and rice leaves. The RAB7^DN^ and RAB7^CA^ mutants failed to produce enough conidia for the pathogenicity assays.

**Figure S4.**
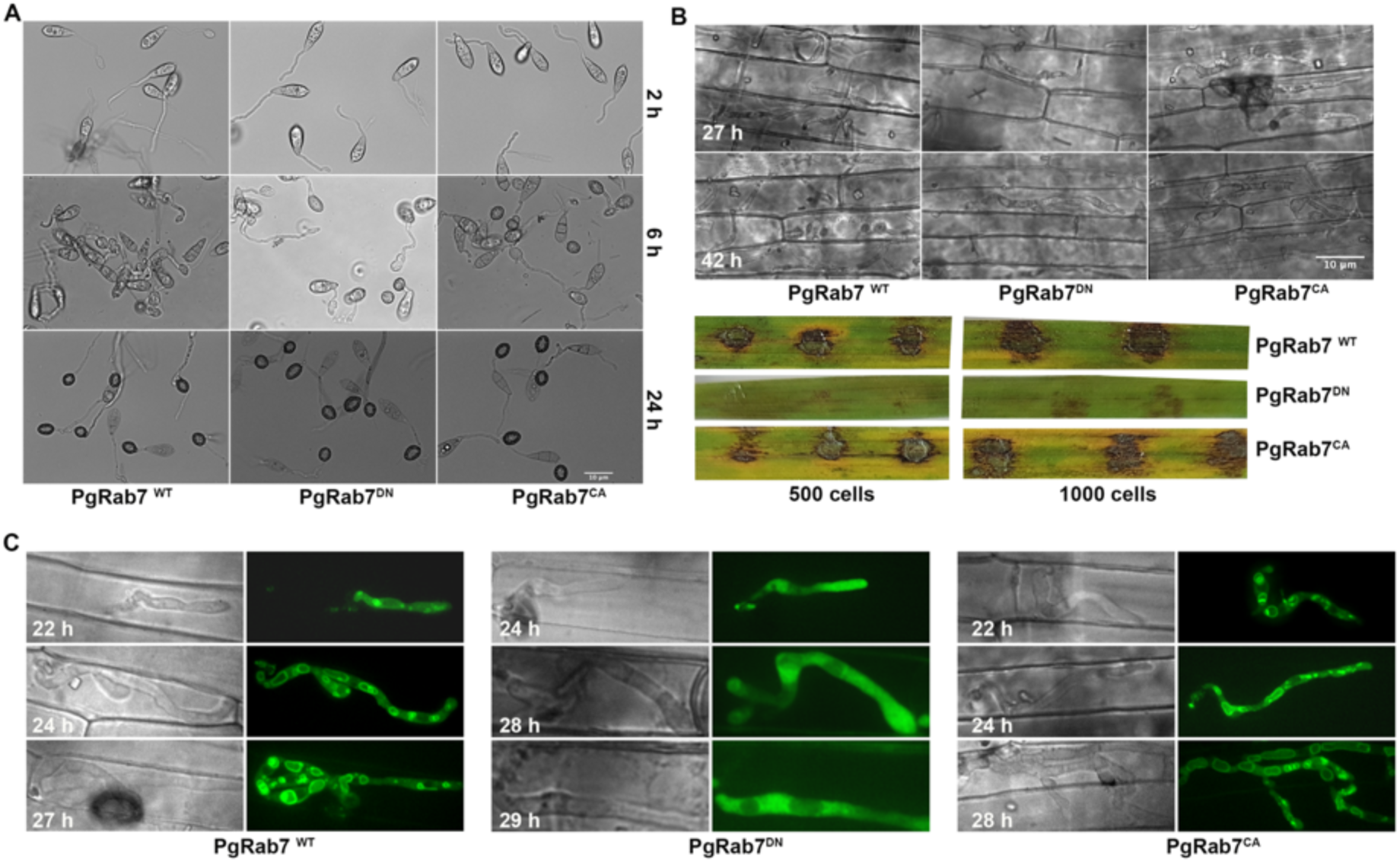
Characterization of the conditional Pwl2-promoter driven *GFP-RAB7* mutants or derivatives that are expressed only in the invasive hyphae. (A) Micrographs showing the conidiation and appressorium formation in the conditional mutant versions of GFP-Rab7. (B) Proliferation of the invasive hyphal growth in rice leaf sheath at indicated time points. The PgRab7^DN^ mutant shows delayed host penetration and is unable to proliferate into the neighbouring cells even at 42 h (upper panel). The mutant is unable to produce significant lesions even at higher inoculum density while PgRab7^CA^blast lesions are comparable to the wild-type PgRab7^WT^ infections (bottom panel). (C) The localization of GFP-Rab7 in WT and mutant backgrounds. Rab7 localizes to late endosomes and vacuole in PgRab7^CA^ mutant comparable to PgRab7^WT^ while it becomes cytosolic in the PgRab7^DN^ mutant, the resultant invasive hyphae of the mutant are unable to proliferate when compared to the WT or the CA mutant.

**Figure S5.**
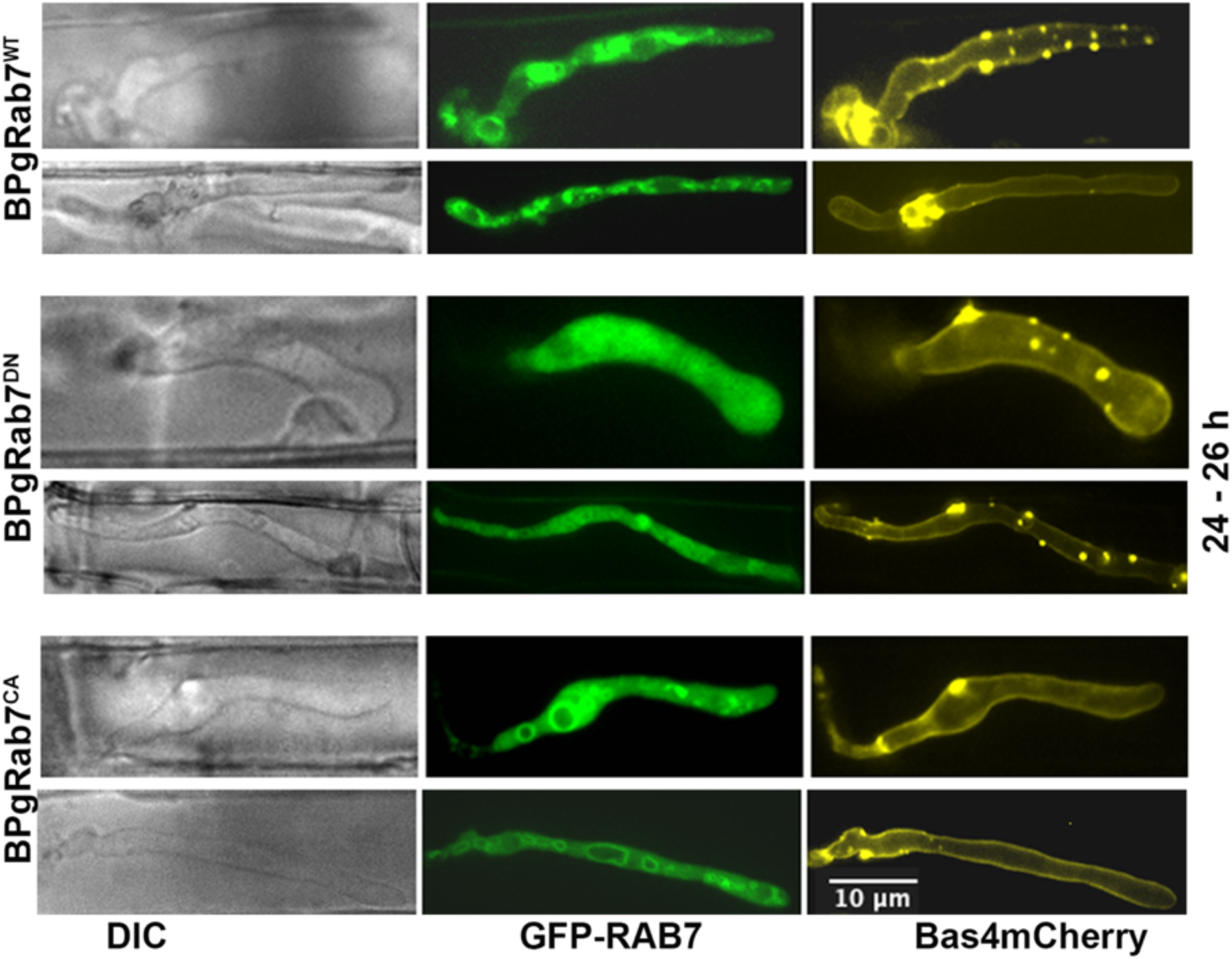
Localization of GFP-RAB7 and Bas4-mCherry during the biotrophic stage in the conditional mutants of RAB7. GFP-RAB7 localizes to cytoplasmic vesicles and vacuolar membrane in BPgRab7^WT^, whereas it becomes completely cytoplasmic in the BPgRab7^DN^. Bas4-mCherry localization remains unperturbed in the BPgRab7^DN^ and BPgRab7^CA^ and comparable to the BPgRab7^WT^ at the indicated timepoints.

**Figure S6.**
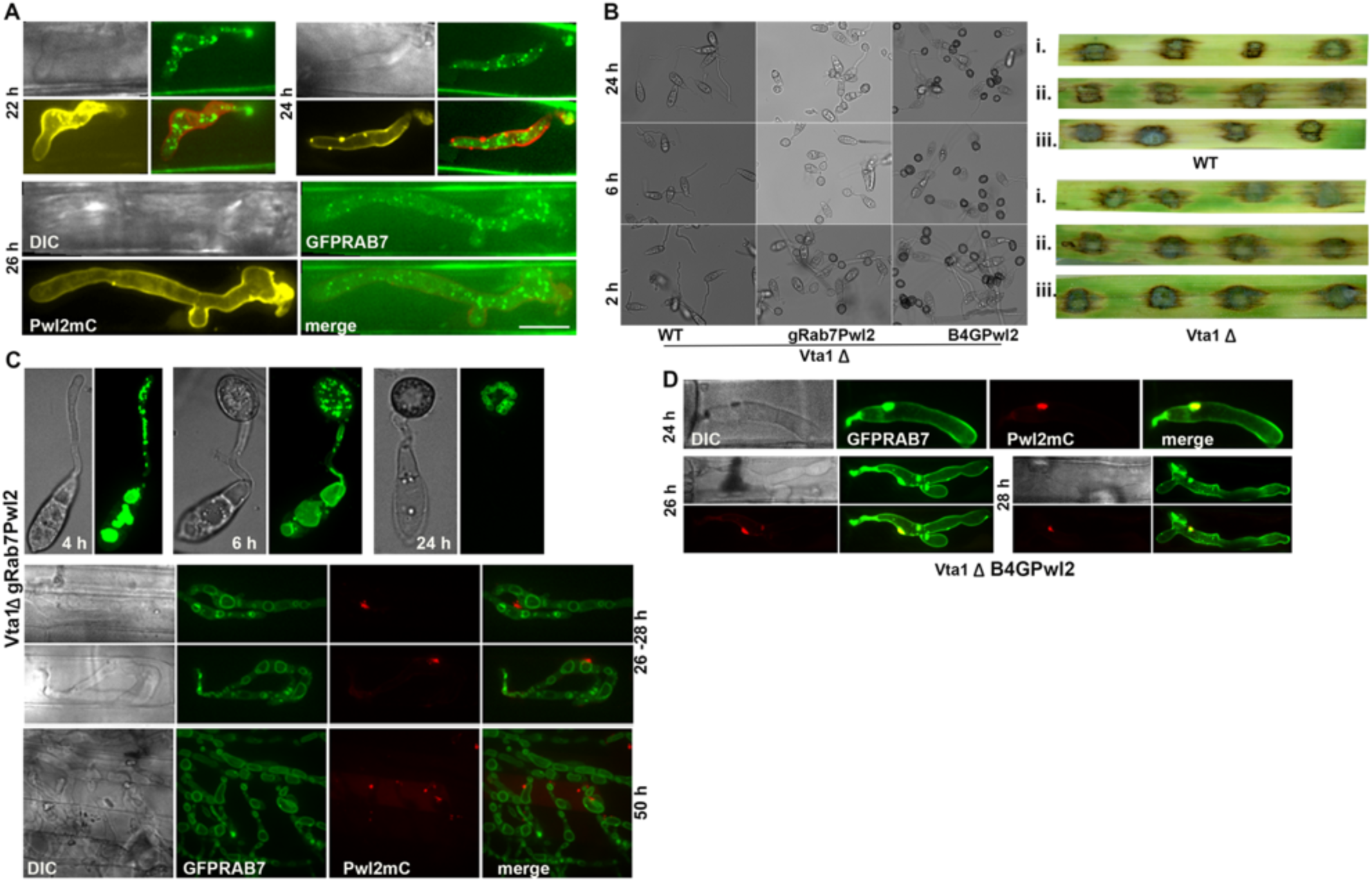
Loss of Vta1 has no effect on Bas4 localization. (A). Vta1-GFP and Bas4-mCherry localization in Vta1gB4mC at different time points during the biotrophic growth. Vta1-GFP punctae do not co-localize with Bas4mC at any instance. (B) Characterization of Vta1 deletion mutant in WT, gRab7Pwl2 and B4GPwl2. Vta1 deletion does not affect the conidiation, appressorium formation or pathogenicity (right panel) of *M. oryzae*. The top panel leaves show the pathogenicity of WT strains (i) WT, (ii) gRab7Pwl2, (iii) B4GPwl2 and the bottom panel the Vta1 deletion in the corresponding strains, respectively. (C) Confocal images showing the localization of GFP-Rab7 and Pwl2-mCherry in the Vta1-deletion strain. The MVB formation seems to be affected while the BIC localization remains unaltered. (D) Localization of Bas4gfp or Pwl2mCherry is unaltered upon Vta1 deletion. The images show the localization of Bas4gfp and Pwl2mC in the invasive hyphae of Vta1ΔB4GPwl2 at the indicated time points of biotrophic growth.

## Supplementary Movies

Videos S1-S3. Time lapse videos of GFP-Rab7 in gRab7 marking the late endocytic events in the pathogenic structures.

Video S1- Localization of GFP-Rab7 at 4h. GFP-Rab7 localized to vacuoles and as vesicles traverse from the conidia to the emerging appressoria as a continuum of tubular vesicles is seen.

Video S2. Localization of GFP-Rab7 at 6h – Rab7 marked vescicular trafficking, involving fusion of vesicles as described in Figure 1 is shown.

Video S3. Localization of GFP-Rab7 in the invasive hyphae during biotrophy

Video S4-S5. Time lapse movie showing that Rab7 marked vesicle like organelle is trafficked from the late endosomal structure located beneath into the BIC marked by Pwl2mC in the strain gRab7Pwl2 at 26 (S4) and 28h (S5).

Video S6. Rab7 trafficking to the BIC from the late endosomal structure located beneath into the BIC in PPgRab7^WT^ expressing Rab7 under the *PWL2pro*.

Video S7 – Rab7 trafficking to the BIC from the late endosomal structure located beneath into the BIC in PPgRab7^CA^ expressing *PWL2pro* PPgRab7^DN^

Video S8 – Time lapse movie showing that Rab7 marked vesicular structures is being trafficked from one organelle to other throughout the invasive hyphae and is rerouted in to BIC when a MVB is under the BIC during biotrophy. The localization of GFP-Rab7 and BIC in gRab7Pwl2 when colocalized with CMAC at 26 h is shown.

Video S9 - A short time lapse showing that it is the membranous counterpart of MVB trafficked to the BIC from the intra lumen of the vacuole that is marked by CMAC.

Video S10 and S11 –Rab7 marked vesicle that is trafficked to the BIC is rich in PI3P marked by GFP-2xFYVE

Video S12 – Vta1-mScarlet vesicle localizes to the membrane of vacuoles or MVBs

Video S13-S15 – Time lapse and surface rendered (S13 and S15) movies showing that Vta1-GFP is highly mobile and is trafficked to the BIC.

## Notes

https://doi.org/10.5281/zenodo.14038222

### Competing Interest Statement

The authors have declared no competing interest.

