## Supplementary Figures for "Phospholipid signaling and multivesicular endolysosomes modulate membrane dynamics at the biotrophic interface in Rice Blast"

**Selvaraj et al 2024: Supplementary Figures**

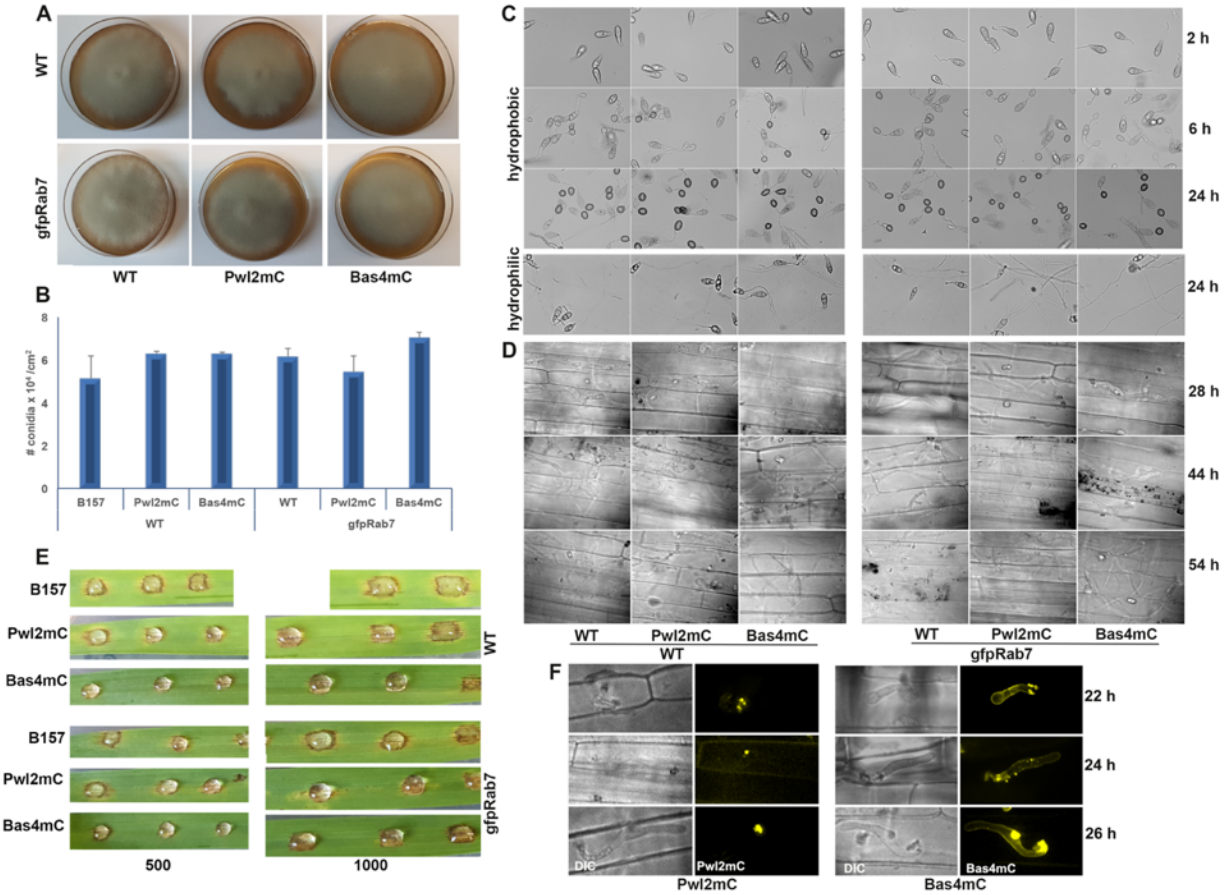


**Figure S1.**

Characterization of strains co-expressing GFP-Rab7 with Pwl2-mCherry (gRabPwl2) or Bas4-mCherry (gRabBas4) representing a cytoplasmic and apoplastic effector, respectively.

(A) Colonies showing the growth of GFP-Rab7 strains expressed in the WT B157 background, Pwl2-mCherry and Bas4-mCherry strains compared with the corresponding WT. (B) Bar graph showing the conidiation levels in the indicated strains. Each column represents mean ± standard error (S.E) of three independent replicates and the experiment was repeated twice.

(C) Images showing the conidial germination and appressorium formation in the indicated strains under hydrophobic or hydrophilic conditions.

(D) Micrographs showing the invasive hyphal growth in rice leaf sheath at indicated timepoints.

(E) Blast disease lesions produced by GFP-Rab7 Pwl2-mCherry or Bas4-mCherry strains compared to the corresponding wild type in barley (top) and rice leaves (bottom).

(F) Confocal images showing the localization of Pwl2-mCherry and Bas4-mCherry in the respective strain co-expressing GFP-Rab7.

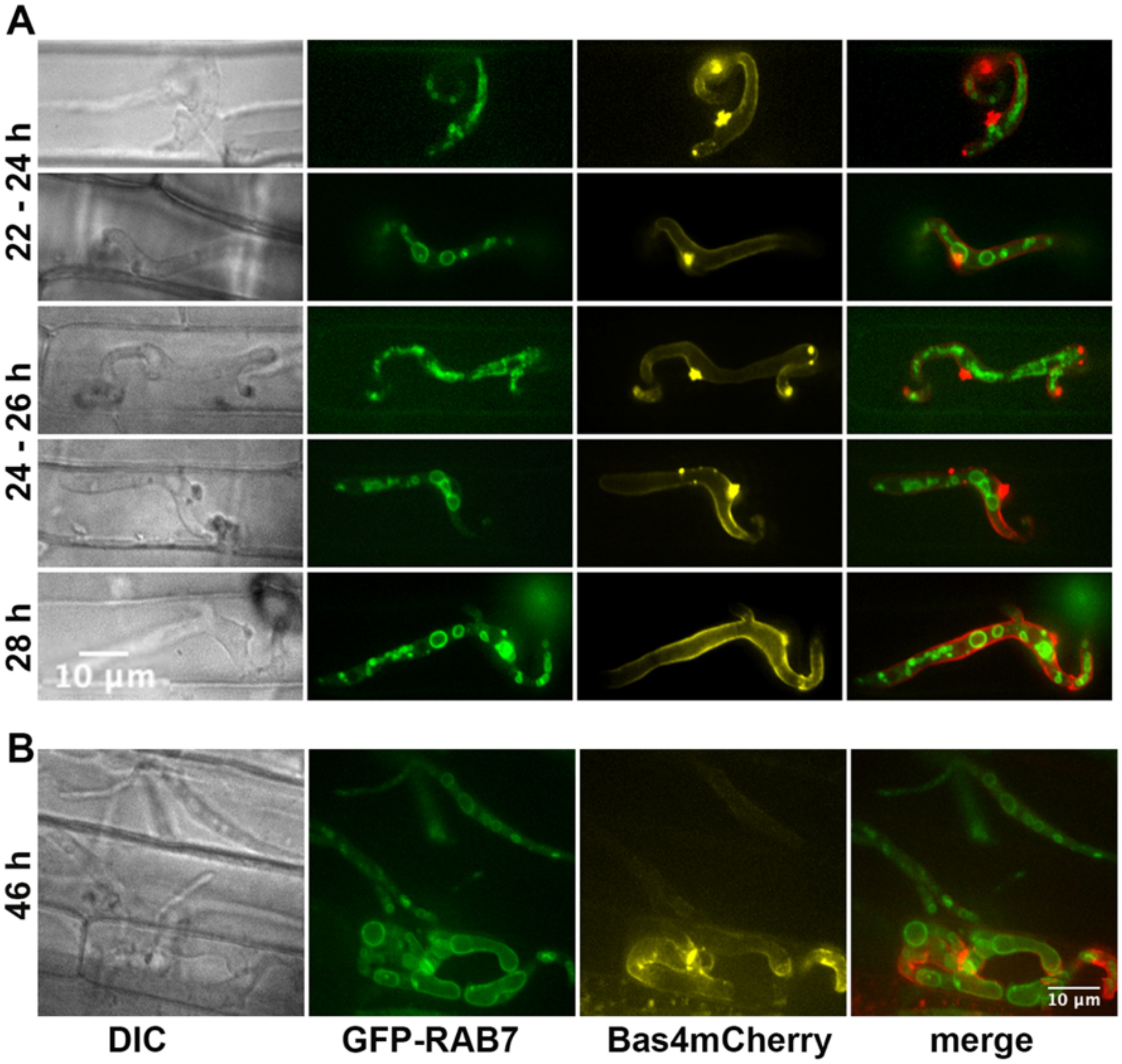


**Figure S2.**

Localization of GFP-Rab7 and Bas4-mCherry in the biotrophic hyphae at (A) earlier time points during the tip, side and focal BIC stages and (B) later time points (46 h) in gRab7Bas4. Bas4-mCherry localizes to the extra-invasive hyphal membrane and GFP-Rab7 localizes to the vesicles and vacuolar membrane but does not co-localize with Bas4-mCherry in the EIHM.

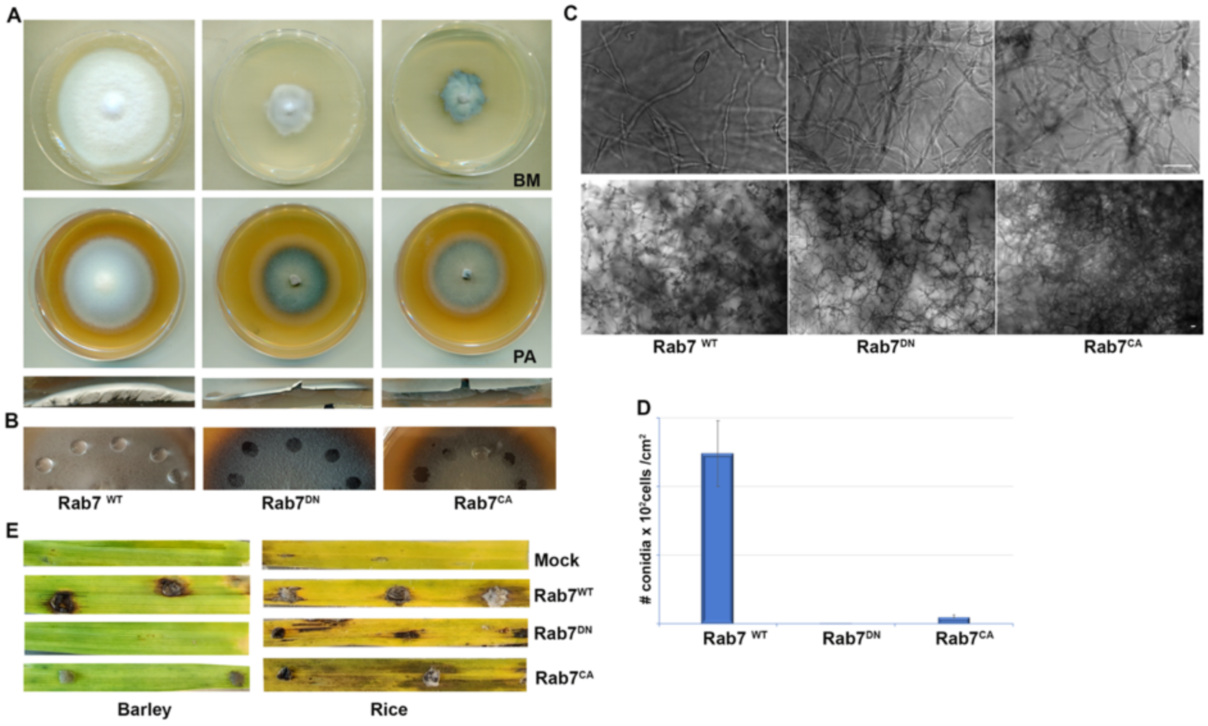


**Figure S3.**

Characterization of Rab7 mutants, the dominant negative (Rab7$^{DN}$) and constitutive active (Rab7$^{CA}$) along with WT (RAB7$^{WT}$).

(A) Colonies showing the radial and aerial growth of Rab7$^{WT}$ or Rab7$^{DN}$, or Rab7$^{CA}$ mutant in Basal medium (BM) or Prune juice agar (PA).

(B) Surface hydrophobicity of Wild type and Rab7 mutants in PA.

(C) Micrographs showing the conidiophore (top) production and conidiation (bottom) in wild type and GFP-Rab7 mutants.

(D) Quantification of conidia produced by WT and the indicated Rab7 mutants. Each value is the mean ± S.E of three individual replicates.

(E) Pathogenicity of the mycelial plugs of WT and GFP-Rab7 mutants in barley and rice leaves. The RAB7$^{DN}$ and RAB7$^{CA}$ mutants failed to produce enough conidia for the pathogenicity assays.

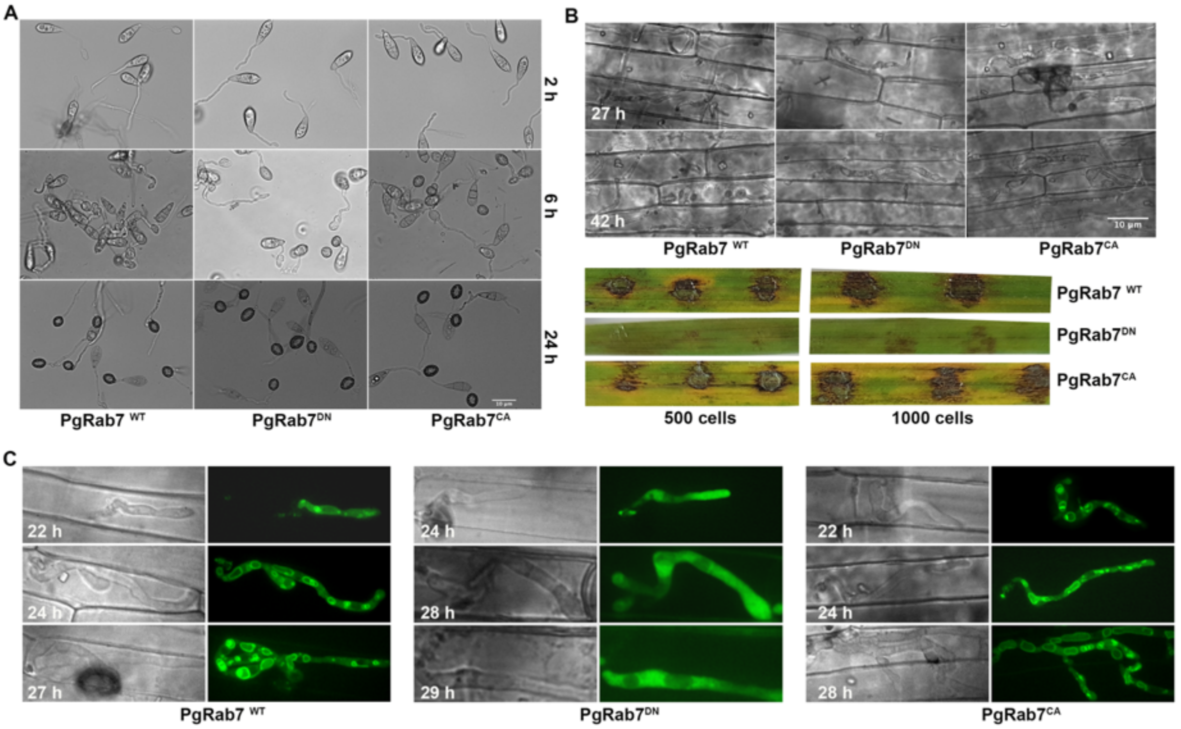


**Figure S4.**

Characterization of the conditional Pwl2-promoter driven *GFP-RAB7* mutants or derivatives that are expressed only in the invasive hyphae.

(A) Micrographs showing the conidiation and appressorium formation in the conditional mutant versions of GFP-Rab7.

(B) Proliferation of the invasive hyphal growth in rice leaf sheath at indicated time points. The PgRab7$^{DN}$ mutant shows delayed host penetration and is unable to proliferate into the neighbouring cells even at 42 h (upper panel). The mutant is unable to produce significant lesions even at higher inoculum density while PgRab7$^{CA}$ blast lesions are comparable to the wild-type PgRab7$^{WT}$ infections (bottom panel).

(C) The localization of GFP-Rab7 in WT and mutant backgrounds. Rab7 localizes to late endosomes and vacuole in PgRab7$^{CA}$ mutant comparable to PgRab7$^{WT}$ while it becomes cytosolic in the PgRab7$^{DN}$ mutant, the resultant invasive hyphae of the mutant are unable to proliferate when compared to the WT or the CA mutant.

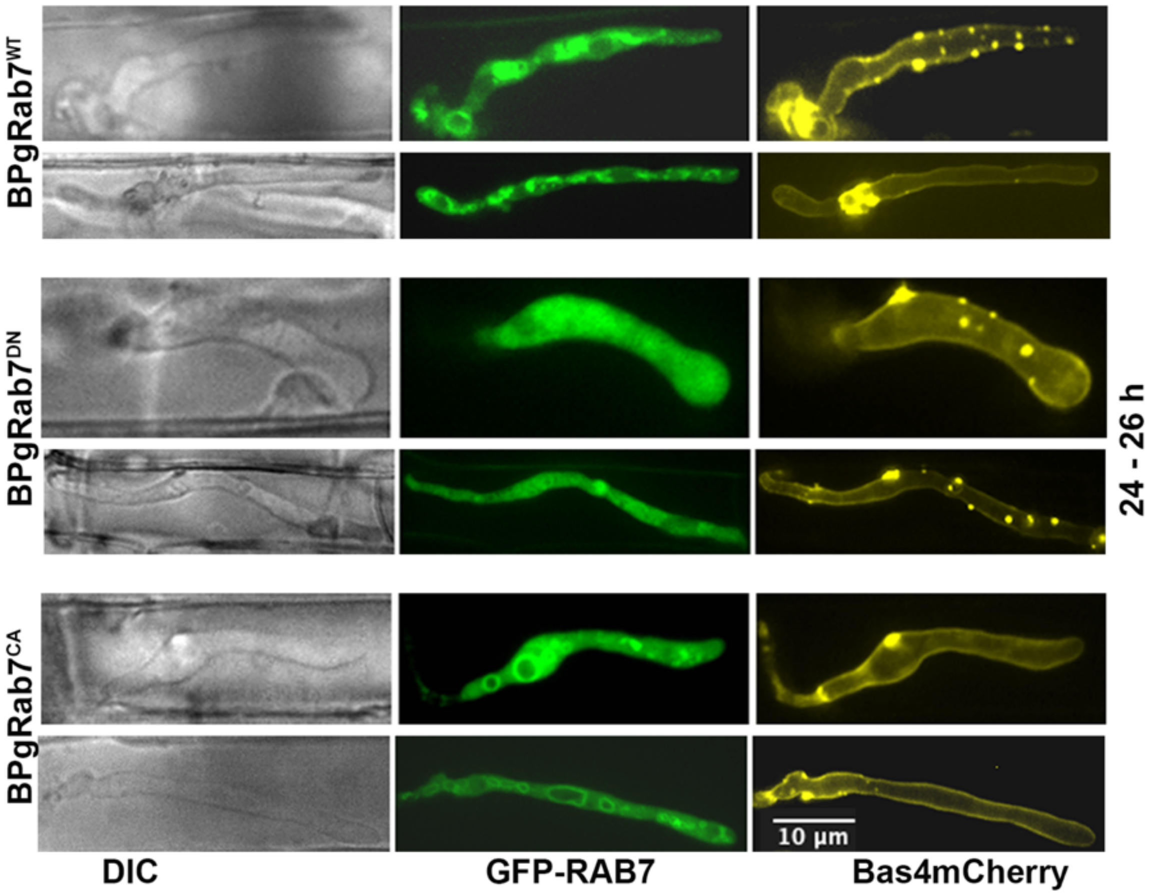


**Figure S5**.

Localization of GFP-RAB7 and Bas4-mCherry during the biotrophic stage in the conditional mutants of RAB7. GFP-RAB7 localizes to cytoplasmic vesicles and vacuolar membrane in BPgRab7$^{WT}$, whereas it becomes completely cytoplasmic in the BPgRab7$^{DN}$. Bas4-mCherry localization remains unperturbed in the BPgRab7$^{DN}$ and BPgRab7$^{CA}$ and comparable to the BPgRab7$^{WT}$ at the indicated timepoints.

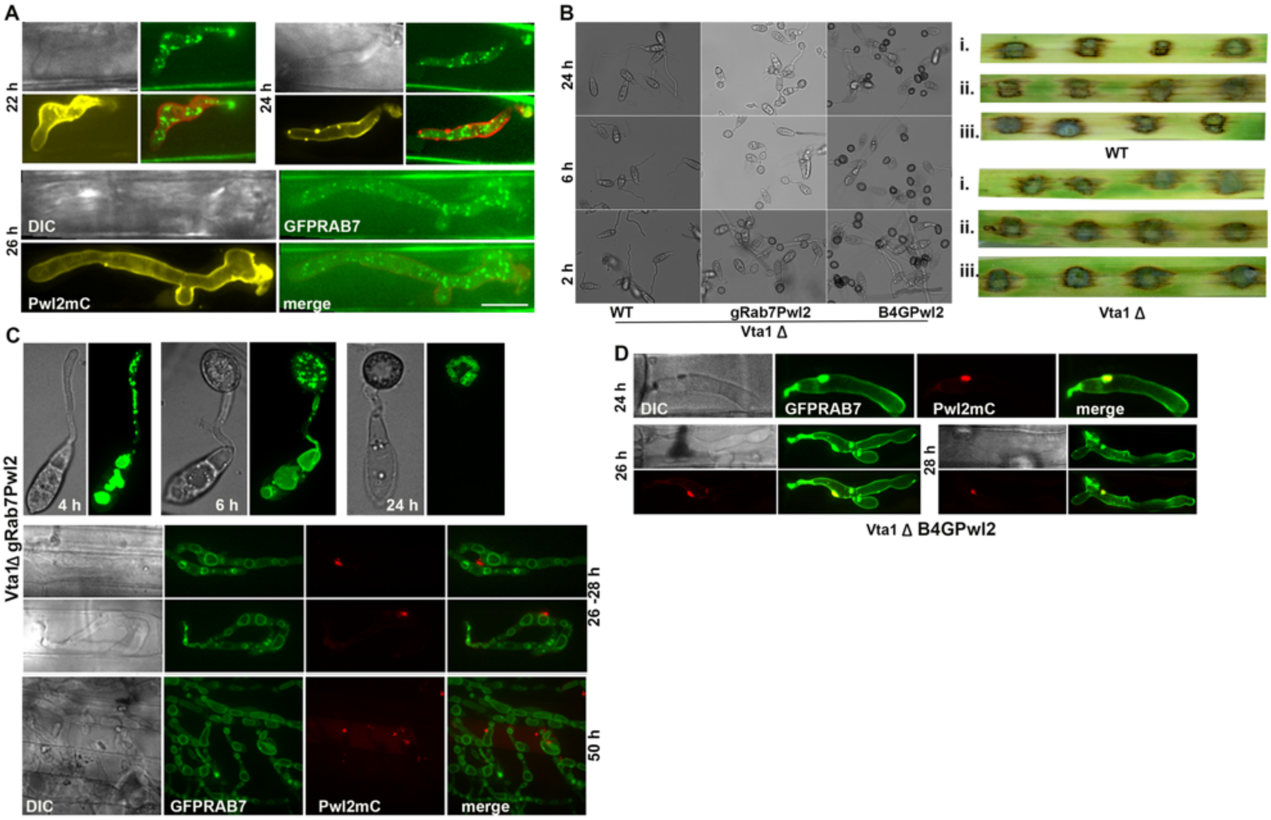


**Figure S6**.

Loss of Vta1 has no effect on Bas4 localization.

(A). Vta1-GFP and Bas4-mCherry localization in Vta1gB4mC at different time points during the biotrophic growth. Vta1-GFP punctae do not co-localize with Bas4mC at any instance.

(B) Characterization of Vta1 deletion mutant in WT, gRab7Pwl2 and B4GPwl2. Vta1 deletion does not affect the conidiation, appressorium formation or pathogenicity (right panel) of *M. oryzae*. The top panel leaves show the pathogenicity of WT strains (i) WT, (ii) gRab7Pwl2, (iii) B4GPwl2 and the bottom panel the Vta1 deletion in the corresponding strains, respectively.

(C) Confocal images showing the localization of GFP-Rab7 and Pwl2-mCherry in the Vta1-deletion strain. The MVB formation seems to be affected while the BIC localization remains unaltered.

(D) Localization of Bas4gfp or Pwl2mCherry is unaltered upon Vta1 deletion. The images show the localization of Bas4gfp and Pwl2mC in the invasive hyphae of Vta1ΔB4GPwl2 at the indicated time points of biotrophic growth.
